# Sensory choices as logistic classification

**DOI:** 10.1101/2024.01.17.576029

**Authors:** Matteo Carandini

## Abstract

Logistic classification is a simple way to make choices based on a set of factors: give each factor a weight, sum the results, and use the sum to set the log odds of a random draw. This operation is known to describe human and animal choices based on value (economic decisions). There is increasing evidence that it also describes choices based on sensory inputs (perceptual decisions), presented across sensory modalities (multisensory integration) and combined with non-sensory factors such as prior probability, expected value, overall motivation, and recent actions. Logistic classification can also capture the effects of brain manipulations such as local inactivations. The brain may implement by thresholding stochastic inputs (as in signal detection theory) acquired over time (as in the drift diffusion model). It is the optimal strategy under certain conditions, and the brain appears to use it as a heuristic in a wider set of conditions.

## Introduction

Many situations require choosing between options based on multiple factors. In a choice between walking and driving, the factors may include distance, traffic, weather, cost, and parking^1^. In a choice between picking a berry vs. moving on, they may include ripeness, hunger, and ease of reach. How does the brain combine such diverse factors to make a choice?

A large body of research indicates that the brain makes choices by logistic classification: perform a weighted sum of the factors, use the result to set the (log) odds of a coin, and toss that coin (Figure 1**a**). If there are more than two options, the coin is replaced by a die. Logistic classification was invented by statisticians in the secrecy surrounding WWII^2^. In the subsequent decades, it was linked to the theory of choice from mathematical psychology and to value-based decisions in economics^1,3,4^. Since then, it has become a workhorse of statistics and machine learning^5^. It characterizes value-based decisions by humans and animals^6-9^ and the corresponding brain activity^7^, suggesting that it was discovered first by evolution.

**Figure 1.**
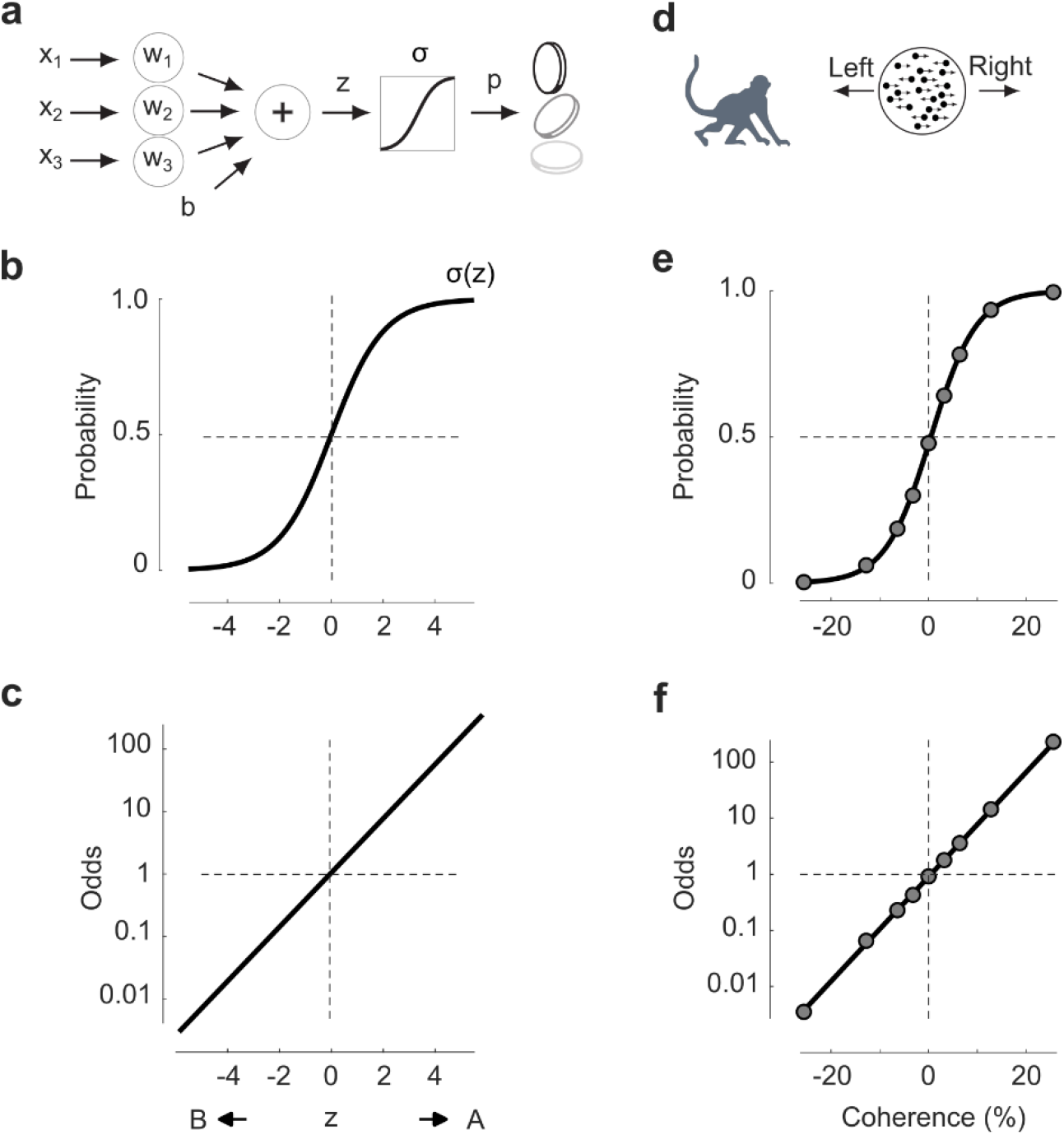
Logistic classification describes sensory choices between two alternatives. **a** Diagram of logistic classification, in the case of 3 factors x_1_, x_2_, and x_3_. The weighted factors are added to a constant bias b and passed through a logistic function. The result is used to bias a coin, which is then flipped. **b**. The logistic sigmoid, plotting probability of choice A vs. decision variable (negative if in favor of choice B, positive if in favor of choice A). **c**. The logistic sigmoid becomes a line when plotting the odds (in logarithmic scale) vs. the decision variable. **d** A monkey performing the random dots task, indicating whether most dots move Left or Right. **e**. Performance of an example monkey, showing probability of Right choices as a function of coherence (negative for leftward motion, positive for right-ward motion). Data were taken from Ref.^11^ using a data grabber (grabit.m). Curve shows fit of the logistic classification model. **f** Same data and fit, in log odds.

There is increasing evidence that logistic classification describes not only choices based on considerations of value (economic decisions) but also choices based on sensory input^2,10,11^ (perceptual decisions) and on additional factors, alone or in combination. These factors include trial history and prior probability^12-22^, disparities in value^7-9^, signals from additional sensory modalities^23^ (multisensory integration), and effects of brain manipulations such as local inactivations^24-26^.

Here I briefly review this evidence, and I argue that logistic classification is an effective description of results that had been previously analyzed with a different framework (signal detection theory^27-29^). I show that it is an optimal strategy in some conditions and a useful heuristic in others, perhaps explaining why it is widely implemented by the brains of multiple species.

## Results

When a choice involves only two alternatives, logistic classification is particularly simple (Figure 1**a**). First, give each of the factors *x*_1_, *x*_2_, … a weight *W*_1_, *W*_2_, … Second, sum them to obtain a *decision variable*

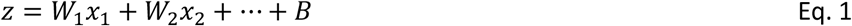

where *B* represents a constant *bias*. Third, use that decision variable to bias a coin, i.e., to set the probability *p* that it will return Heads. Specifically, use it as the *log odds* of drawing Heads:

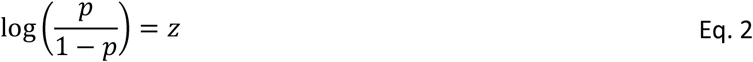

Fourth, flip the coin.

This process can be written as a single equation (Figure 1**a**):

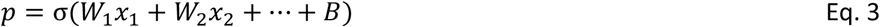

where *σ* is the *logistic sigmoid* (Figure 1**b**)

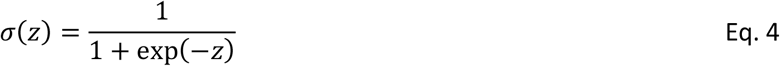

In statistics, these equations are often used for “logistic regression”: to find the weights *W*_1_, *W*_2_, … and bias *B* that best relate the factors *x*_1_, *x*_2_, … to some observed outcomes^3,5^. We are here interested in the reverse: using the weights as given, to classify a new observation of the factors and thus make a choice. This is best called “logistic classification”^5^.

In this description of logistic classification, a choice is stochastic (flipping a coin) and the factors *x*_i_ are deterministic. We will later see an equivalent description where the factors are stochastic and the choice is deterministic, as in signal detection theory^27-29^, which is particularly apt when the factors *x*_i_ are sensory (Section S1). We will also see a description where the stochastic factors are acquired over time, as in the drift diffusion model^2,30-32^ (Section S2). For most of the paper, however, this simple description will suffice.

### Sensory choices between two alternatives

Though it is simple, logistic classification can do an excellent job at summarizing perceptual decisions. For example, in a classic motion discrimination task, a monkey views dots and estimates whether most of them move Left or Right (Figure 1**d**). The task difficulty depends on the coherence *x* of the dots, defined as positive if most dots move to the Right and negative otherwise. The *psychometric curve*, i.e., the probability of Right choices as a function of stimulus strength can then be fit with a logistic classification model (Eq. 3) with a single factor^10,11^:

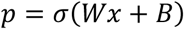

With two free parameters (*W* and *B*) the model describes how nine values of coherence *x* determine the probability of choosing Right (Figure 1**e**).

The model and the data are easier to interpret if instead of probability we plot the logarithm of the odds (Figure 1**f**). In this representation (as shown in Eq. 1), the logistic classification model becomes a straight line:

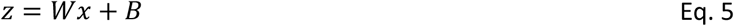

and the measured odds fall neatly on the line (Figure 1**f**). In the rest of the paper, I will thus express the model in terms of *z*.

In this example, the measure of stimulus strength *x* in Eq. 5 is the one used in the laboratory: the coherence of the dots^10,11^. It is often necessary, however, to allow an expansive or compressive deformation that converts the measure used in the laboratory to a scale used inside the brain. For instance, one could raise the stimulus strength to an exponent *q*, which is fit to the data^25,33,34^. For simplicity, I will write this as *x*^*q*^, but the proper expression is *sign*(*x*)|*x*|^*q*^, which maintains the sign of *x*. In other cases, one may want functions *f*(*x*) that saturate more strongly^12,24^.

As we will see later, the weights *W* for sensory factors are bounded by sensory reliability (the inverse of its standard deviation). By contrast, no bound exists for the weights for other factors, such as those based on value. So, one could give an infinite weight to chocolate – ensuring a deterministic choice of dessert – but not to the rightward motion of random dots.

### Sensory choices influenced by priors

In the previous example task (Figure 1**e**,**f**), the stimuli were equally likely to move left or right. In other words, their prior probabilities were equal: *P*_*R*_ = *P*_*L*_. Reflecting this symmetry, the monkey was unbiased (*B* ≈ 0): when the stimulus was uninformative (zero coherence), it responded *L* or *R* with approximately equal probability.

In general, however, one of the choices may have a higher chance of being correct than the other. In this case, the optimal strategy is to set the bias *B* to the logarithm of the prior odds:

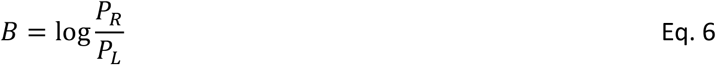

As proven in Section S3, this value of *B* is optimal in that it maximizes the overall reward (if the rewards for the two sides are equal, and if the evidence *Wx* is distributed uniformly).

Therefore, an observer that performs logistic classification and seeks to maximize reward should adjust its bias *B* according to the prior probability of the stimuli (Eq. 6), without changing its sensitivity *W*. The bias may not hit precisely the ideal level (Eq. 6), because the observer may not be able to estimate the true prior probabilities, but it should at least move in the right direction.

This prediction is verified in humans performing a biased motion discrimination task: if the odds of a stimulus going rightward *P*_*R*_ /*P*_*L*_ are 80:20 instead of 50:50 (Figure 2**a**), the observers make more Right choices^35^ especially at zero coherence, when the stimulus is uninformative (Figure 2**b**). This shift is captured by the logistic classification model (Eq. 5) with the same weight *W* but a larger bias *B* (Figure 2**b**). The log odds of the choices fall on a line that is parallel but displaced from the 50:50 case (Figure 2**c**).

**Figure 2.**
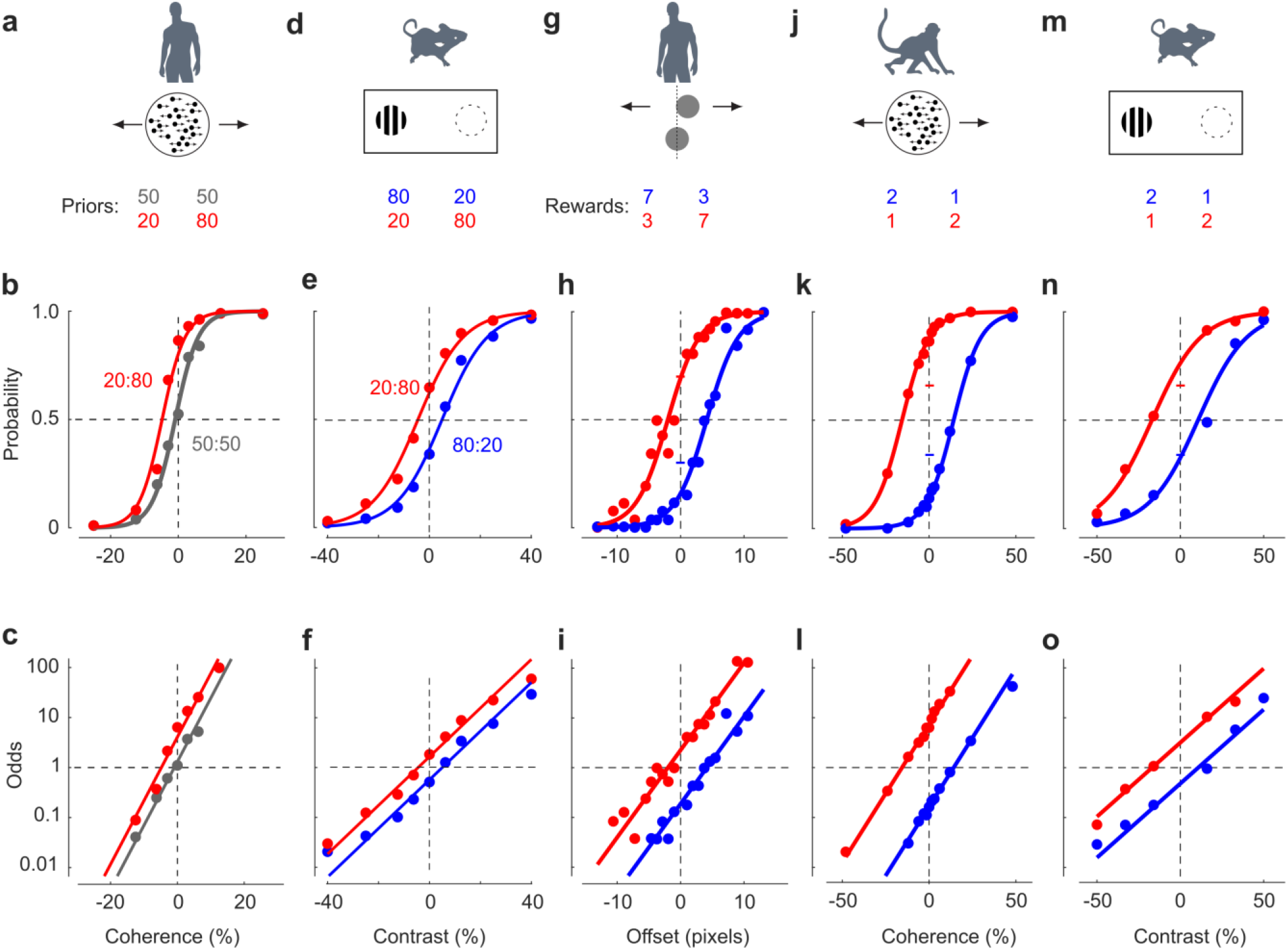
Logistic classification describes choices based on stimuli, priors, and values. **a**. A random dots task where a human is told that rightward motion is more likely than leftward motion (blue) or that they are equally likely (gray). **b**. Data from an observer showing that probability of choosing Right depends not only on stimulus coherence (abscissa) but also on prior probability (blue vs. gray). Data grabbed from Ref.^35^. Curves show fit of the logistic classification model with same weight w but two different values for the bias b. **c**. The same data and curves plotted as odds of choosing Right vs. Left, in logarithmic scale. **d-f**. Same, for a mouse indicating whether a visual stimulus appears on the Left or the Right, presented in alternating blocks where Left was more likely (red) or less likely (blue) than Right. Data grabbed from Ref.^22^. **g**. A position discrimination task where the observer indicates whether the top stimulus is to the Right of the bottom one, in conditions where correct Left responses were rewarded less (red) or more (blue) than correct Right responses. **h**. Data from an observer showing that probability of choosing Right depends not only on stimulus offset (abscissa) but also on reward condition (blue vs. red). Data grabbed from Ref.^45^. Curves show fit of the logistic classification model with same weight W but two different values for the bias B. The red and blue ticks in the central axis indicate the intercepts predicted by the optimal strategy. **i**. The same data and curves plotted as odds of choosing Right vs. Left, in logarithmic scale. **j-l**. Same, for a monkey performing a random dots task where stimuli are presented in alternating blocks where correct Left choices were rewarded less (red) or more (blue) than correct Right choices. Data grabbed from Ref.^46^. **m-o**. Same, for a mouse indicating whether a visual stimulus appears on the Left or the Right, presented in alternating blocks where correct Left choices were rewarded less (red) or more (blue) than correct Right choices. Data grabbed from Ref.^47^.

Analogous results are seen in mice that perform a biased position discrimination task^22^ (Figure 2**d**). In this experiment, stimuli are divided in blocks having higher probability on the Left (80:20) or on the Right (20:80). As expected, the psychometric curves measured in the two blocks are displaced (Figure 2**e**,**f**), such that at zero contrast (no visual stimulus) Right choices are more likely in 20:80 blocks and Left choices more likely in 80:20 blocks. The model (Eq. 5) captures this difference with a bias *B* that is positive in the 20:80 block and negative in the 80:20 block, with no changes in sensory weight *W*.

### Sensory choices without evidence

In the absence of sensory evidence, these data exhibit a phenomenon that is widely observed in non-sensory tasks^8,9,36-43^, called “probability matching”: the probabilities of the observer’s choices tend to match the probabilities of those choices being correct. For instance, at zero coherence the human in the 20:80 motion task chose Right with probability ∼0.8 (Figure 2**b**). At zero contrast, the mouse in the position task showed a similar (though less pronounced^9,44^) tendency (Figure 2**e**).

This phenomenon is puzzling because it is suboptimal^37,40^. Indeed, in the absence of evidence, one would obtain more rewards by always guessing Right if *P*_*R*_ > *P*_*L*_ and always guessing Left otherwise (Section S3). In other words, one should set *B* to +∞ if *P*_*R*_ > *P*_*L*_ and to −∞ otherwise (in practice, setting *B* to +10 or -10 would suffice).

The observer, however, does not know that some trials have no evidence. It may thus apply the optimal value of the bias (Eq. 6) to all trials. This strategy produces probability matching. Indeed, if the evidence is uninformative (*x* = 0), then the probability that the observer chooses *R* is

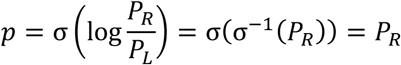

where I used the fact that log 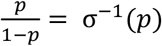.

Perhaps brains use Eq. 6 as a heuristic (even when they should be setting *B* = ±∞), because in most situations in nature there is a correct answer that can be inferred from factors (present stimuli, past stimuli, actions, outcomes, etc.). This strategy is suboptimal when no factors are relevant (e.g. when gambling), and leads them to perform probability matching.

Reassuringly, humans stop performing probability matching if they understand that they are gambling (guessing an independent random variable). In that case, they become more optimal, privileging the option that is more likely to be rewarded^40^.

We have seen that if an observer performs logistic classification (Eq. 5) they should set their bias to the log prior odds (Eq. 6), which leads to probability matching (when the evidence is zero). This argument can also be inverted: as shown in Section S4, the only way to achieve probability matching (for any strength of evidence) is to perform logistic classification (Eq. 5) and to set the bias to the log prior odds (Eq. 6).

### Sensory choices influenced by value

In the experiments examined up to here, the rewards for correct Left and Right choices were equal. In general, however, the rewards, *V*_*L*_ and *V*_*R*_, could differ. In this general case, one can prove (Section S3) that the ideal bias *B* (the bias that maximizes reward) is

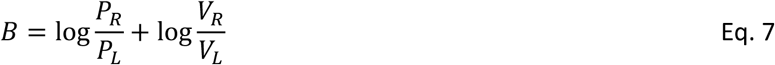

Therefore, if the probabilities of both options are equal (*P*_*R*_ = *P*_*L*_) an optimal logistic classifier should engage in “value matching”: its fraction of Right choices should match the ratio of Right rewards to total rewards. Indeed, if *x* = 0 the probabilities of Right choices should be *p* = σ(log(*V*_*R*_ /*V*_*L*_)) = *V*_*R*_ /(*V*_*R*_ + *V*_*L*_).

There is ample evidence that the brain follows this strategy when making purely economic decisions^7^, even though it would obtain more reward by sticking to the option with higher value. Once again, logistic classification suggests an explanation for this behavior: the brain may assume that there is relevant evidence *x* to guide the choice (even when there cannot be any), so it may obey Eq. 7 even when not optimal.

More generally, the optimal logistic classifier (Eq. 7) captures how observers combine sensory signals with considerations of value. For example, when humans perform a position discrimination task (Figure 2**g**), they make more Left or more Right choices depending on which choice yields the larger reward^45^. The data are well fit by logistic classification (Eq. 5) with a single sensory gain *W* and different biases *B* in the two reward conditions (Figure 2**h**). Once again, the effects are easiest to judge in terms of log odds (Figure 2**i**): changing the stimulus moves the log odds along a line, and changing the reward ratio moves the log odds to a parallel line. Similar results are seen in monkeys doing the random dot motion task^46^ (Figure 2**j-l**), and in mice performing the position discrimination task^47^ (Figure 2**m-o**). In all these cases the changes in bias were at least as large as predicted by optimality (Eq. 7). In fact, they were generally larger than optimal^48^, perhaps because of a non-linear utility scale that disproportionally favors large rewards^7,49^.

Taken together, logistic classification (Eq. 5) and the ideal bias (Eq. 7) indicate how to combine sensory signals with relative probability and relative reward. They show that the logarithm of the odds provides a “natural currency for trading off sensory information, prior probability and expected value”^10^.

### Sensory choices influenced by motivation

In the experiments examined up to here, observers assigned a constant weight *W* to sensory evidence (resulting in a constant slope of the log odds line) in the face of changes in stimulus strength, probability, and value. However, paying attention to the evidence may require motivation. If observers are more motivated, they may give a higher weight *W* to the evidence.

For instance, in an experiment measuring rate discrimination in rats, changes in the ratio of rewards were yoked to changes in the total available reward^26^. Logistic classification provides good fits to those data (Supp Figure 1) and reveals changes both in bias *B* and in sensory weight *W*. The changes in bias are expected given the changes in ratio of rewards (Eq. 7). The changes in sensory weight, instead, might result from the changes in total available reward, which might affect task engagement^12^ or sensory attention^50^. Variations in these factors may explain why the parameters of logistic classification often fluctuate over time^51,52^.

### Sensory choices across modalities

Logistic classification can describe not only sensory choices based on a single modality but also those based on multiple modalities (multisensory integration). As shown in Eq. 1, logistic classification naturally accommodates multiple factors as input (Figure 1**a**). If there are two sensory stimuli *s*_1_, *s*_2_ of different modalities (e.g., vision and audition), and these stimuli are statistically independent, Bayes’ rule (as shown in Section S4) predicts that an observer should perform logistic classification based on their sum^23^:

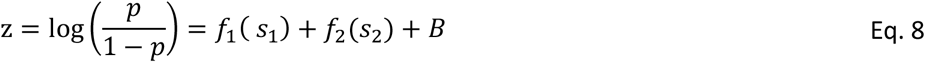

This expression is just the sum of three factors, and yet it describes multisensory choices in a variety of species, tasks, and combinations of modalities.

For instance, Eq. 8 describes the choices of mice performing an audiovisual detection task^23^. In this task, a visual pattern appears on the Left or on the Right and a sound is played in one of 3 or 5 positions (Figure 3**a**). If the sound is in the middle, it is uninformative, and the task becomes purely visual (Figure 3**b**, *gray*). If it is on the Left or Right, the mouse is more likely to make those choices (Figure 3**b**, *blue* and *red*).

**Figure 3.**
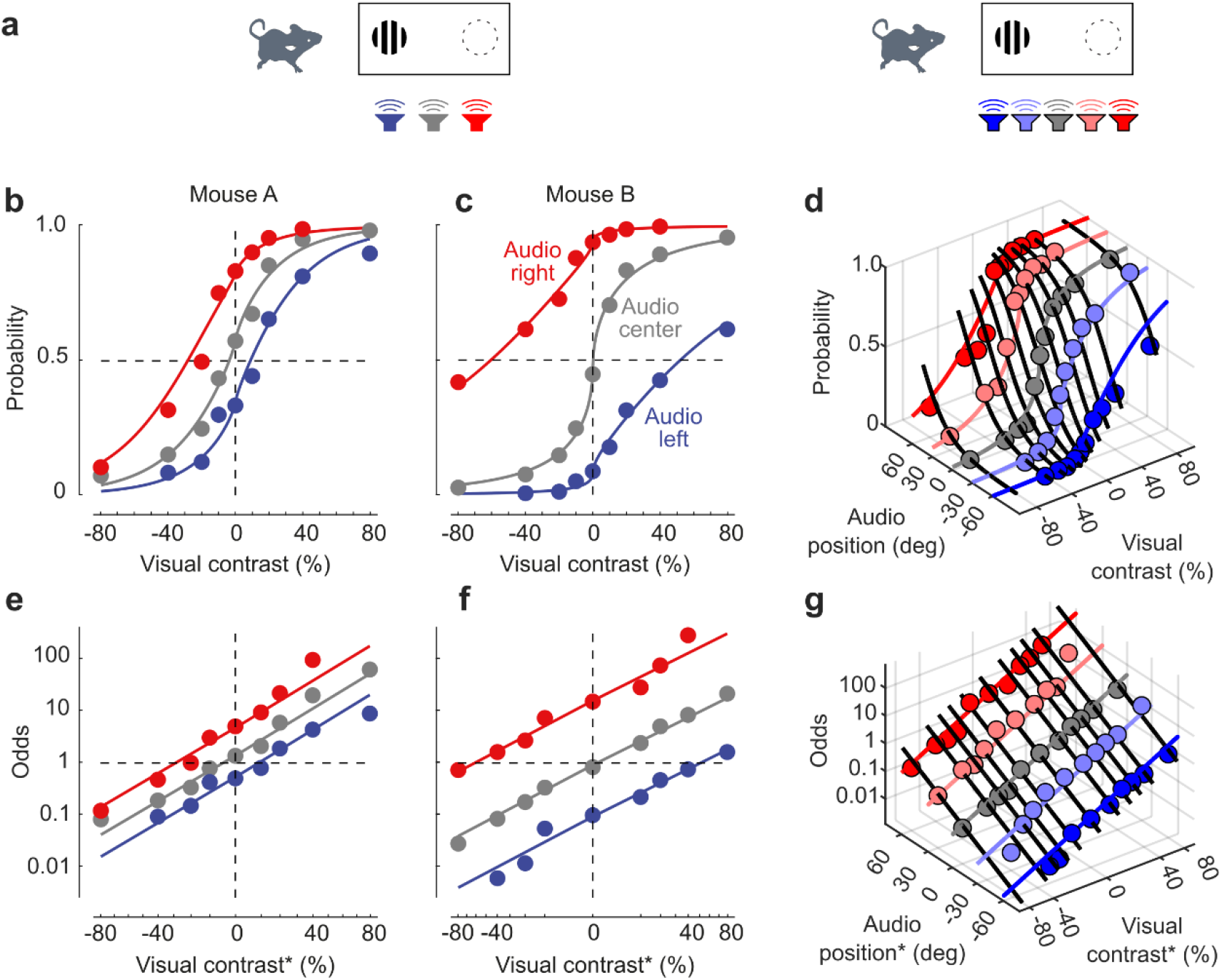
Logistic classification describes audiovisual choices. **a**. The audiovisual task: images appear on the Left or right, and sounds are emitted at one of 3-5 positions. **b**. Probability that a mouse chooses Right based on the contrast of the visual stimulus (abscissa, negative for stimuli on the Left) and on the position of the auditory stimulus (colors as in a). **c**. Same, for a mouse with higher auditory sensitivity. **d**. Results in a task with 5 auditory positions. **e-g**. The data in b-d, replotted as odds of Right vs. Left choices, in logarithmic scale. The quantities on the abscissa (horizontal plane in g) are compressed (raised to an exponent q<1). Data from Ref.^23^. Therefore, the probability of choosing R is a logistic function p = σ(z) with three terms: one that depends on vision, one that depends on audition, and a constant.

Logistic classification (Eq. 8) describes this behavior accurately, as the sum of three factors:

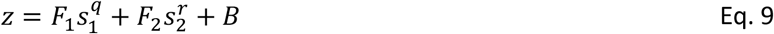

Here, the variables *s*_1_ and *s*_2_ are visual and auditory attributes, raised to exponents *q, r* (if necessary). The remaining parameters are the visual gain *F*_1_, the auditory gain *F*_2_, and the bias *B*. The first term describes the responses when the task is purely visual (Figure 3**b**, *gray*), and the second describes the responses when the task is purely auditory (Figure 3**b**, where abscissa is zero). The model provides excellent fits (Figure 3**b**, *curves*), correctly describing the choices not only when the two modalities agree, but also when they contradict each other^23^. Similar results are seen in a mouse with better auditory localization (higher performance at zero contrast, Figure 3**c**), and in a mouse tested with more auditory positions (Figure 3**d**).

As usual, these results are easiest to understand in terms of log odds (Figure 3**e**-**g**). In this representation, and with a slightly different abscissa (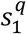 and 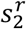 instead of *s*_1_ and *s*_2_) the results fall on parallel lines: changing visual contrast moves the points along the lines, and changing the auditory position shifts them to a different line.

For such high-dimensional data, one way to judge the performance of the model^31,53^ is to plot the probability of all points as a function of the predicted decision variable *z* (Eq. 1) and see if they fall near the logistic curve *σ*(*z*) (Eq. 4). Here they do (Supp Figure 3).

Logistic classification describes multisensory choices in multiple other species across multiple modalities. Studies of multisensory integration have typically analyzed the responses with signal detection theory (Section S1), and found that observers are roughly^54,55^ Bayesian^23,26,56-61^ (at least when all stimuli are presented for the same duration^62^): they weigh each modality according to its reliability^56,57^. Below I examine three such studies^55,59,61^, which tested multisensory integration with different tasks, across different modalities (visual, auditory, tactile, and vestibular), in different species (rats, monkeys, and humans), and I show that their data are well described by logistic classification.

In a study that measured visuo-tactile integration^55^, rats indicated whether the orientation of a stimulus was closer to horizontal or vertical by choosing between Left or Right response ports. The stimulus could be purely visual or purely tactile (Supp Figure 2**a**,**b**) or a combination of the two (Supp Figure 2**c**). I applied the logistic classification model to those data with 4 free parameters: visual gain *F*_1_, tactile gain *F*_2_, their common exponent *q* (there was no need for different exponents), and bias *B*. The model provided good fits (Supp Figure 2**a**-**c**), and would have done fairly well even with just two free parameters (setting *q* = 1 and *B* = 0, not shown). The model’s performance is easy to evaluate if the data are plotted as log odds: aside from the (small) bias, the line for multisensory stimuli (Supp Figure 2**f**) is just the sum of the lines for unisensory stimuli (Supp Figure 2**d**,**e**). An additional check verifies that the predicted decision variables for all these measured data describe a single logistic curve (Supp Figure 3**a-c**).

### Sensory choices for multimodal features

The other two studies^59,61^ tested multisensory integration in more detail, by varying both the amplitudes and the features of two modalities, and by introducing offsets between those features.

To discuss these experiments, it helps to define stimulus “feature” vs. “amplitude”. The feature *ϕ* is the attribute on which the choice is based; it can be positive or negative, and is zero at the boundary between the two choices. The amplitude *α*, instead, is positive and is zero if the stimulus is absent. For instance, in the motion task (Figure 1**d**-**f** and Figure 2**a**-**c**), *ϕ* is direction (±1) and *α* is coherence (between 0 and 1). Similarly, in the position task (Figure 2**d**-**f**), *ϕ* is position and *α* is contrast.

If a stimulus *s* can have multiple combinations of amplitude and feature, we must define how those combinations affect the stimulus representation *f*(*s*). A reasonable working hypothesis is that the stimulus representation is separable, i.e., the product of a function of amplitude and one of feature: *f*(*ϕ, a*) = *g*(*ϕ*)*h*(*α*). In the visual system, for example, this hypothesis would be appropriate for orientation and contrast^63^ (but perhaps not for temporal frequency and contrast^64^).

Under this hypothesis, when there are two sensory modalities and thus two amplitudes *α*_1_, *α*_2_ and two features *ϕ*_1_, *ϕ*_2_, the model (Eq. 8) becomes

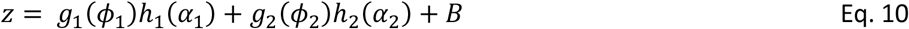

To test this model, I considered a task where humans received sequences of visual and auditory stimuli and judged whether their rate was above a criterion^60,61^. As expected, when stimuli were unisensory, the observers did well if the stimulus had high amplitude (“Easy”), and poorly if it had low amplitude (“Hard”, Figure 4**a**,**b**). When the easy auditory stimulus was combined with the hard visual stimulus, it largely determined the observer’s choices: if the two stimuli disagreed, the observer responded in a manner consistent with the auditory stimulus (Figure 4**c**). Similar results were obtained by combining the hard auditory stimulus with the easy visual stimulus (Figure 4**d**). In both cases the choices were largely determined by the easier stimulus (as expected from a Bayesian observer^56,57,61^).

**Figure 4.**
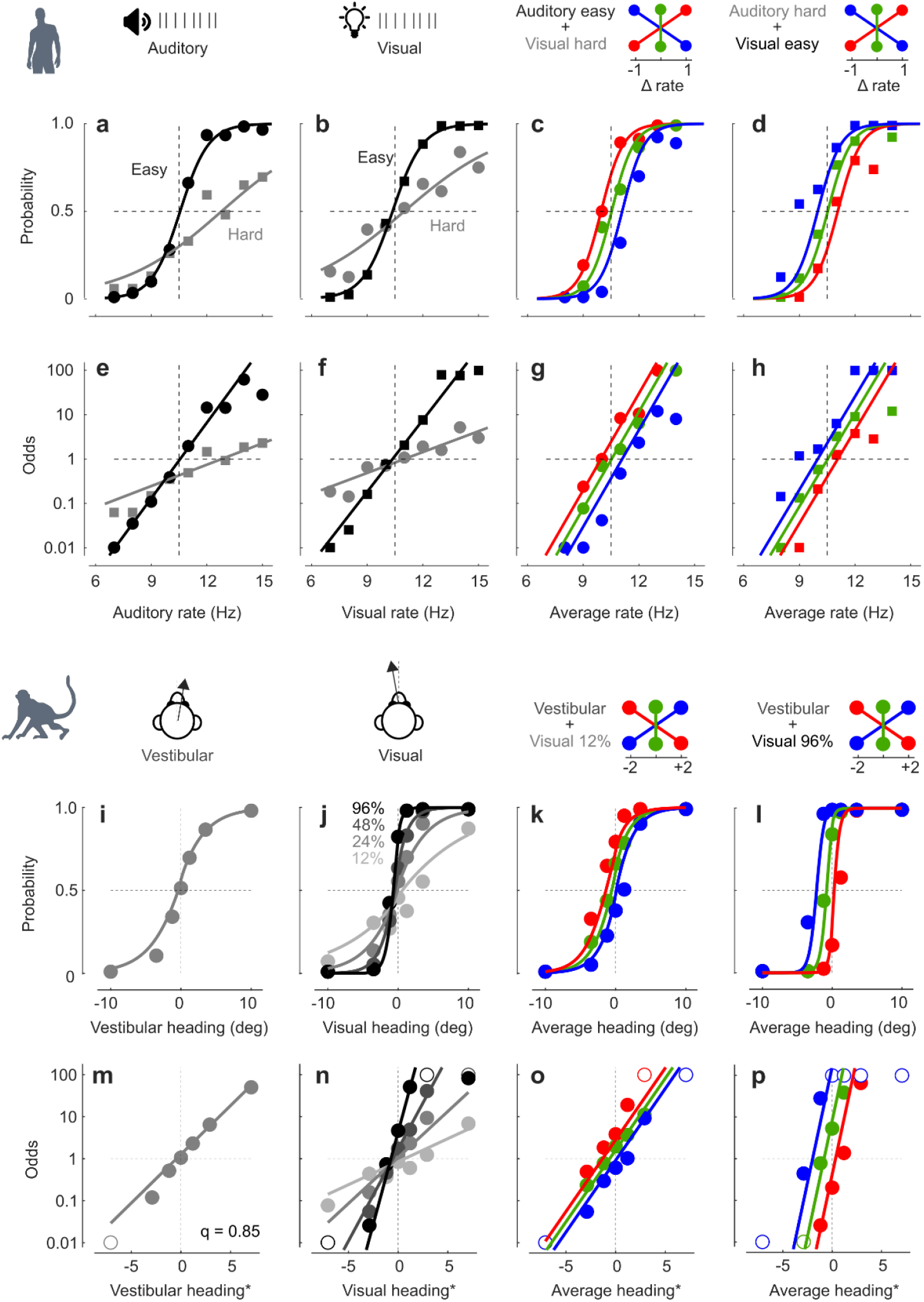
Logistic classification describes audiovisual and vestibulo-visual choices with stimuli differing in feature and amplitude. **a** Average choices of a human performing a rate discrimination task based on auditory sequences. The ordinate plots the probability that the observer judges the rate of the sequence to be higher than a reference (10.5 Hz) when stimuli have low amplitude (Hard) or high amplitude (Easy). Curves in this panel and subsequent ones are fits of the logistic classification model. **b**. Same, for visual stimuli. **c**. Same, for stimuli obtained by summing an easy auditory stimulus to a hard visual stimulus. As illustrated in the inset, the rate of the auditory stimulus was lower (blue), equal (green), or higher (red) than the rate of the visual stimulus. The abscissa plots the average rate of the visual and auditory stimuli. **d**. Same, for stimuli obtained by summing a hard auditory stimulus to an easy visual stimulus. **e**-**h:** The same data as a-d, plotted in terms of odds, in logarithmic scale. Data are from Human 1 in Ref.^61^. **i** Average choices of a monkey performing a heading discrimination task based on unisensory vestibular stimulation. The ordinate plots the probability that the observer judges the heading to be Right. Curves are predictions of the logistic classification model. **j**. Same, for unisensory visual stimuli varying in coherence from 12% (hardest) to 96% (easiest). **k**. Same, for multisensory stimuli obtained by summing the vestibular stimulus to a hard visual stimulus, with the former having a heading left of (blue), same as (green), or right of (red) the latter. The abscissa plots the average heading of the vestibular and visual stimuli. **l**. Same, for stimuli obtained by summing the vestibular stimulus to an easy visual stimulus. **m**-**p:** The same data as a-d, plotted in terms of odds, in logarithmic scale. Open symbols indicate values where the probability is close to 0 or 1, where the odds would require a high precision, and are thus likely to lie outside the range of the ordinate. Data were grabbed from Ref.^59^.

The choices measured in this task agree with logistic classification, with amplitudes and features combined under the separable approximation (Eq. 10). To fit the data, I chose linear expressions for the functions: *h*_*i*_(*α*_*i*_) = *α*_*i*_, and *g*_*i*_(*ϕ*_*i*_) = *F*_*i*_ *ϕ*_*i*_ + *B*_*i*_ (with *i* = 1,2 denoting visual and auditory modalities). The result is the following expression:

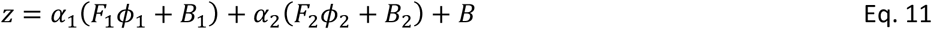

With just 5 free parameters (visual gain *F*_1_, visual bias *B*_1_, auditory gain *F*_2_, auditory bias *B*_2_, and overall bias *B*) this function produced 10 psychometric curves that fitted 114 measurements (Figure 4**a**-**d**, *curves*).

As usual, model and data are easiest to inspect as log odds. In this scale, the unisensory responses are lines that are steeper for Easy stimuli than for Hard stimuli (Figure 4**e**,**f**). When the steep line from one modality is summed to the shallow line from the other, its largely determines the slope and intercept of the resulting line (Figure 4**g**,**h**).

In another multisensory task, monkeys judged whether their heading was Left or Right based on visual and vestibular cues^59^. The animals performed well when presented with purely vestibular stimuli (Figure 4**i**) and with purely visual stimuli with high coherence (Figure 4**j**). When the vestibular stimulus was combined with the hardest visual stimulus (lowest coherence), the former largely determined the observer’s choices (Figure 4**k**). Conversely, when the vestibular stimulus was combined with the easy visual stimulus, the latter largely determined the choices (Figure 4**l**). As expected from a Bayesian observer^59^, therefore, the choices were largely determined by the modality that was easier.

These data were well fit by the logistic classification model under the separable assumption (Eq. 11). Here, *ϕ*_1_indicates the visual heading and *ϕ*_2_ indicates the vestibular heading, both raised to an exponent *q* to capture increased sensitivity around the heading of 0. This function provided good fits to the psychometric curves (Figure 4**i**-**l**), and as usual, the reasons for its success are easiest to see in terms of log odds (Figure 4**m**-**o**).

It is striking that logistic classification describes multisensory choices in so many species, tasks, and modalities, especially considering that the tasks do not conform to the conditions where logistic classification is optimal.

Indeed, logistic classification (Eq. 8) is optimal if the modalities are independent^23^ (Section S4), but here the two modalities had positive correlations. For instance, in the audiovisual position task (Figure 3) the modalities agreed more often than they disagreed^23^. More strikingly, in the visuotactile task (Supp Figure 2) the two modalities had a correlation of 1 when both present, because they always had the same orientation. Strong correlations between modalities were present also in the other two tasks (Figure 4). In these conditions, logistic classification may no longer be optimal, but the brain seems to adopt it nonetheless as a useful heuristic^65^.

### Sensory choices influenced by history

A key challenge in making a choice is to weigh only the factors that are relevant: giving weight to irrelevant factors decreases the chance of correct answers. And yet, superstitious behavior is widespread. For instance, in sensory tasks where the sequence of stimuli is random, humans and other animals^12-22,49,66-75^ typically act as if it followed some order (“serial dependence”).

The logistic classification model can capture this serial dependence by including history factors as regressors in addition to the sensory factors^12-22,73,76^. This approach was obtained to explain apparently erratic choices exhibited by mice in a visual task^12^. It merged previous logistic models that had separately used sensory factors^10^ and history factors^7-9^.

In this framework (Figure 5**a**), choices depend not only on the present sensory stimulus *x*(*t*) as in Eq. 5, but also on whether the preceding choice was a success *s* or a failure *f*:

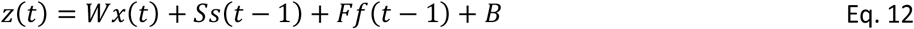

where *s*(*t*) = *sign*(*x*(*t*)), *f*(*t*) = 0 if trial *t* was a success, and *s*(*t*) = 0, *f*(*t*) = −*sign*(*x*(*t*)) if it was a failure.

**Figure 5.**
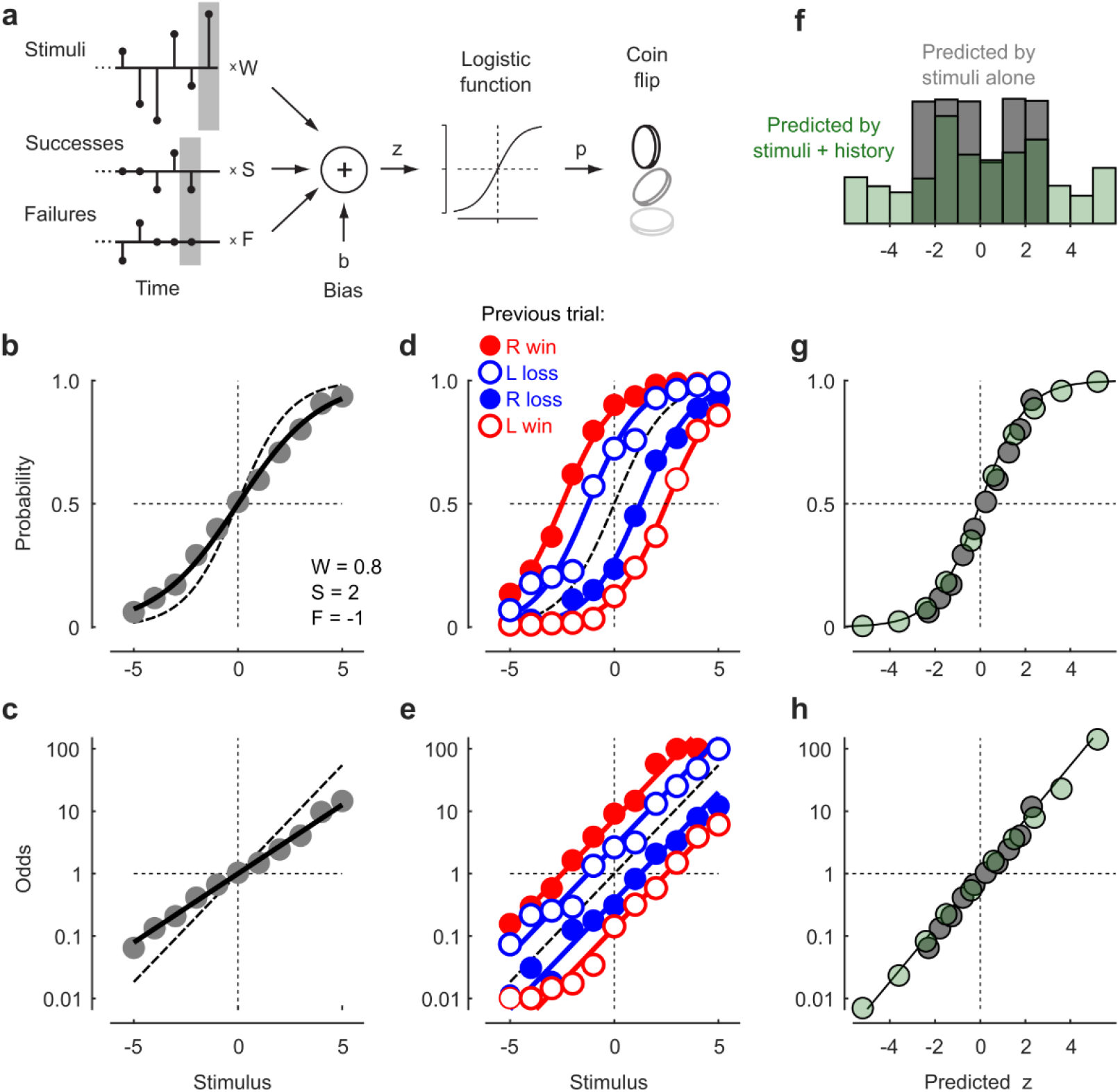
Logistic classification describes effects of history. **a**. Model of logistic classification where choices are determined not only by a stimulus but also by prior wins or losses (Eq. 12). Adapted from Ref.^12^. The subsequent panels show results of simulations with W = 0.8, S = 2, F = -1, B = 0. See Fig. 2c of Ref. ^74^ for similar data. **b**. Simulated data showing the probability of R choices as a function of contrast, compared to what would be observed based on the sensory factor alone (dashed). **c**. Same, plotted as the logarithm of the odds. **d-e**. Same as b-c, contingent on previous trial being a success or a failure (red vs. blue) on the L or R (open vs. closed). The dashed curve shows the result that would be observed based on the sensory factor alone. **f**. Distribution of values of the decision variable z, as predicted by the stimuli alone (gray) or by the stimuli plus the history terms (green). **g**. Binned values of the decision variables for the two models, on top of a logistic curve. **h**. Same, plotted as the logarithm of the odds.

This equation describes just one of various possibilities, chosen for its simplicity. In the studies that used this approach^12-22,73,76^, the history regressors typically extend further in the past, or are expressed in different combinations such as past choices (regardless of success) or past rewards (regardless of choice).

If serial dependence is present, ignoring it can make one underestimate stimulus sensitivity. Indeed, plotting the performance as a function of the present stimulus yields shallower psychometric curves (fewer correct answers) than would be observed based only on sensory sensitivity *W* (Figure 5**b**,**c**). A better estimate of sensitivity (and a measure of the effect of history) is obtained by plotting performance conditional not only on the present stimulus but also on the choice and outcome of the previous trial (Figure 5**d**). If the effect is additive (as in Eq. 12), it will simply shift the lines representing the log odds of the choices, because the history factors sum with the fixed bias *B* to yield a variable overall bias (Figure 5**e**). One can confirm that these history factors are in play by comparing a model that ignores them (Eq. 5) to one that includes them^53^ (Eq. 12). In our example, the model that includes history factors predicts larger (positive or negative) values for the decision variable *z* (Figure 5**f**), while providing good fits (Figure 5**g**,**h**).

The strength of the history terms varies across individuals and in time^12^. It can change with learning^51^ and can be increased or decreased by appropriate design, but it cannot be removed^16^. In humans, for instance, it correlates with education^16^: PhD candidates or graduates have strongly negative failure weights (lose-switch) but weak success weights, whereas the remaining observers have positive success weights (win-stay). This correlation is not necessarily causal and may reflect other unmeasured factors.

However, some effects of history are more complex than can be captured by a simple logistic classification model such as Eq. 12. For instance, past choices have a stronger effect if they were made with low confidence^71^, a behavior that is more readily explained by models based on reinforcement learning^47^. Also, engagement in the trial typically fluctuates, and these fluctuations can be captured by allowing the weights to change across trials^51,52^. Variations in engagement will vary the fraction of successes, so observing data after a success vs. a failure may be a way to gauge the effect of engagement. This phenomenon may explain results observed in rats during rate discrimination^26^, where past successes and failures affect not only the overall bias but also the sensory weight *W*, which was slightly higher after a success than after a failure (Supp Figure 1**g**-**l**). Given that successes tend to occur in an engaged state and failures in a disengaged state, the latter effect may be explained by a dependence of sensitivity on engagement.

### Sensory choices during brain manipulations

Logistic classification is a good model not only for how the brain makes decisions, but also for how these decisions are modified by brain manipulations such as local inactivations^23-26^. If the effects of inactivating a brain region can be summarized by changes in a few weights, one can surmise that the region is necessary for processing the corresponding factors.

For example, in mice performing the audiovisual localization task (same as Figure 3), inactivation of the right visual cortex decreases the contralateral visual weight, but no other weights. As a result, it impairs the detection of visual stimuli on the left side, but does not affect visual detection on the right side or auditory detection on either side (Figure 6**a**). This effect is particularly easy to see in terms of log odds (Figure 6**b**).

**Figure 6.**
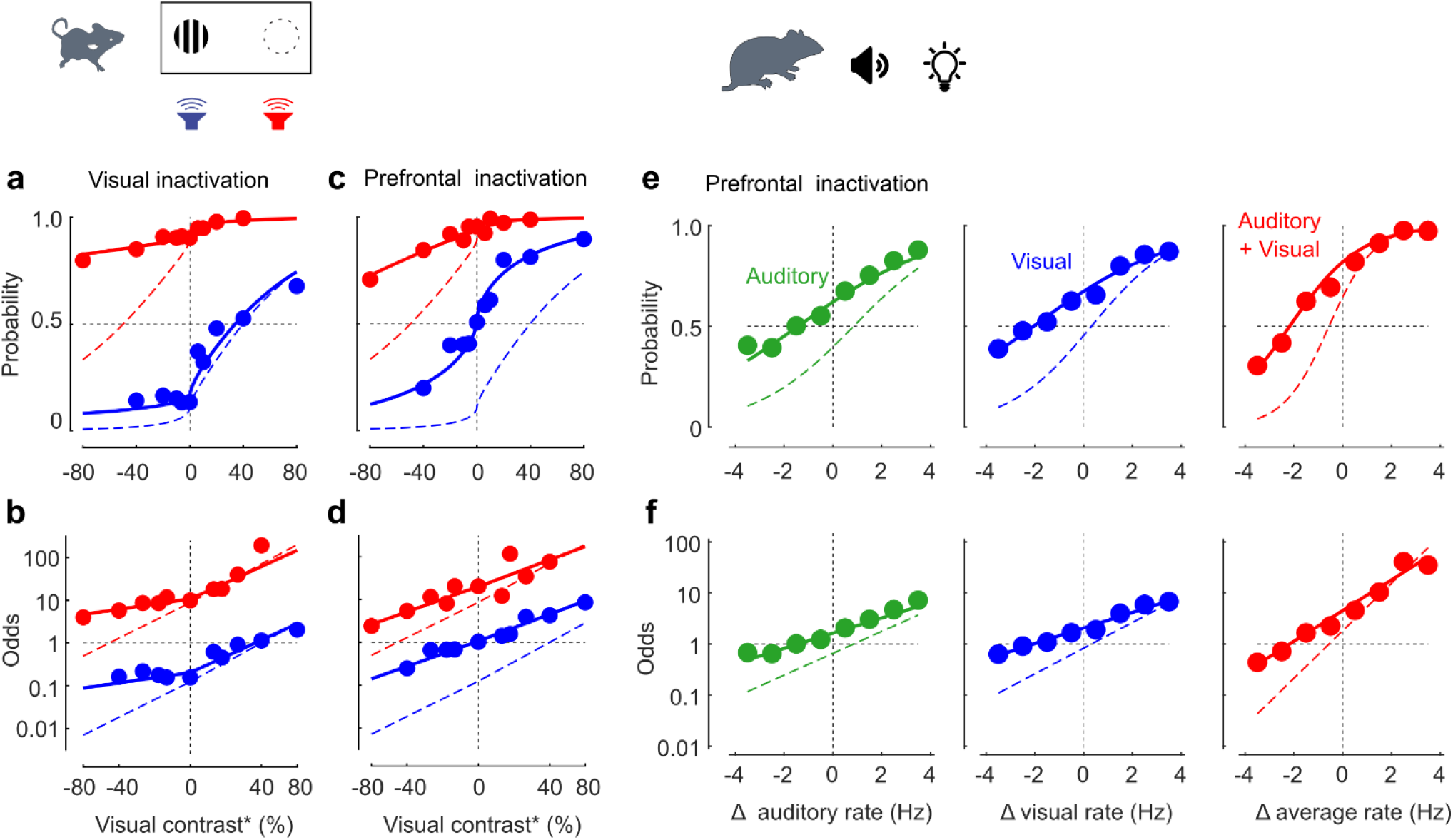
Logistic classification describes effects of brain manipulations. **a**.Effects of inactivating right visual cortex in mice that perform an audiovisual localization task^23^. Showing the probability of Right choices as a function of visual contrast (abscissa) in the presence of sounds from the Left (blue) and Right (red), in control conditions (dashed) and during inactivation (data and continuous curves). The curves are predictions of the model (Eq. 14) allowing only one parameter to change with inactivation: the weight applied to visual stimuli on the left, F_1 L_. **b**. Same data, plotted as log odds. **c**,**d**. Effects of inactivating the right prefrontal cortex. The model captured them via changes not only in the contralateral visual weight but also in the contralateral auditory weight and in the overall bias. Replotted from Ref. ^23^. **e**. Effects of prefrontal inactivation in a rat performing rate discrimination based on auditory (green), visual (blue) and audiovisual (red) cues, showing the probability of choosing “Right” (for high rates) as a function of rate (expressed relative to the reference). **f**. Same data, in log odds. Data from Ref. ^26^.

To describe these asymmetric effects, it helps to extend Eq. 9 to allow for different weights for stimuli on the left vs. right^23^. To this end, we split the decision variable *z* into two variables *z*_*L*_, *z*_*R*_ summarizing the evidence in favor of the two choices:

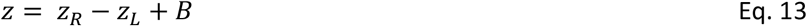

and we give each of them a visual weight (*F*_*L*,1_ and *F*_*R*,1_) and an auditory weight (*F*_*L*,2_ and *F*_*R*,2_):

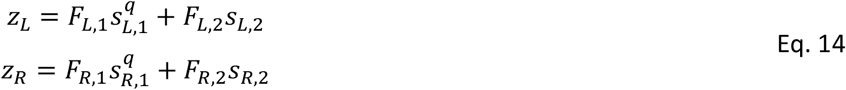

This model has 6 parameters (four sensory weights, the exponent, and the bias), and in principle a brain manipulation could change all those parameters relative to the control condition. Typically, however, the control condition is described by fewer parameters (e.g., the weights on the left and right are the same), and brain manipulations change only a few parameters^23-26^. For instance, inactivating right visual cortex affects only the weight applied to left visual stimuli, *F*_*L*,1_ (Figure 6**a**,**b**). This result indicates that right visual cortex is only necessary for processing visual stimuli, and only those on the left^23^.

By contrast, inactivating right prefrontal cortex reduces both visual weights and auditory weights (regardless of side), and increases bias *B* thus increasing the proportion of Right choices (Figure 6**c**,**d**). This result indicates that prefrontal cortex processes bilateral visual and auditory signals and promotes contralateral choices^23^.

Similar results are seen in rats performing the audiovisual rate discrimination task^26^ (same as Figure 4): the control and inactivation conditions are well fit by logistic classification (Eq. 14), and inactivation reduces both auditory and visual weights (regardless of whether stimuli have low or high rate), and increases the bias (Figure 6**e**-**j**).

### Sensory choices without flipping coins

These results indicate that it is useful to think of the brain as computing separate variables *z*_*L*_, *z*_*R*_ summarizing the evidence in favor, and more generally the utility, of each choice (Eq. 13). Each of the variables is obtained by a weighted sum of factors (similar to Eq. 1). The overall decision variable *z* is obtained by subtracting the utilities of the two choices (Figure 7**a**).

**Figure 7.**
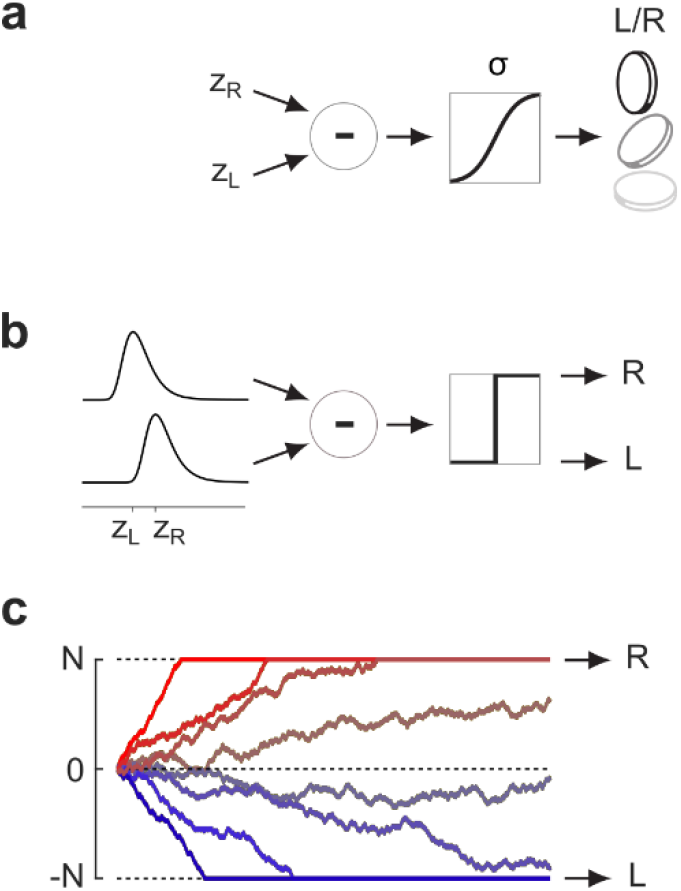
Three implementations of logistic classification. **a**. The implementation described in most of this paper, based on deterministic utilities z_L_, z_R_ informing a stochastic decision. **b**. An alternative implementation where the two utilities are stochastic across trials, and are then subtracted and thresholded. This is analogous to signal detection theory^27-29^. **c**. An implementation where the decision variable z controls the drift of a random walk, and the choice is made when the walk hits one of two boundaries. This is the widely used drift diffusion model^2,30-32^.

However, this implementation still involves a set of deterministic factors followed by a stochastic decision (a coin flip). The brain may be faced with the opposite situation: a set of stochastic factors, especially if these factors are sensory, followed by a deterministic choice. Logistic classification can be achieved also this way, in alternative implementations that result in the same equations used throughout this paper.

The first alternative implementation has strong similarities to signal detection theory^27-29^ (Figure 7**b**). In this implementation, the two utilities *z*_*L*_, *z*_*R*_ determine the means of two random variables, which vary from trial to trial (e.g. because of sensory noise) and are distributed according to a specific distribution (called a Gumbel distribution). The two utilities are then subtracted, and if the result is above zero the choice is R, otherwise it is L. This approach is used in economics^1,3^ to model differences among consumers, and in signal detection theory^27-29^ to model sensory noise (Section S1). In this implementation, the weight *W* that an ideal observer would assign to a sensory factor (Eq. 1) is proportional to the factor’s reliability (the inverse of standard deviation of the noise). This represents an upper limit: a less ideal observer would assign a lower weight.

The second alternative implementation is analogous to the widely used drift diffusion model^2,30-32^. In this implementation, a single decision variable *z* controls the drift of a random walk, and the choice is made when the walk hits one of two boundaries (Figure 7**c**). This implementation is identical to the logistic classification model if the starting point is midway between the two bounds^31,77^. Indeed, as detailed in Section S2, if it takes *N* steps to reach a bound^78^, and the probability to go up (vs. down) at each step is set to *P* = *σ*(*z*/*N*), then the probability of hitting the upper bound is *p* = *σ*(*z*). If one chooses *z* as the weighted sum of factors (Eq. 1), this is the logistic classification model.

The drift diffusion model predicts not only the probability but also the timing of the choices^2,30-32^ (Section S2). For instance, it predicts that larger values of a decision variable result in faster decisions.

There are many ways to change the drift diffusion model, e.g. with an asymmetric starting point or with diverse relationships between *P* and sensory inputs and biases^79^. In some of those cases there may not be a closed form equation for the probability of hitting a boundary. In those cases, the psychometric functions must be computed numerically^80^.

### Sensory choices between more alternatives

In all the analyses above, we have assumed that the observer faces a binary choice. However, this is not a strict requirement: The logistic classifier is readily extended to the case when the choice is one of *n* alternatives. First, one obtains a vector of decision variables ***z*** = *z*_1_, *z*_2_, …, *z*_*n*_, one for each alternative:

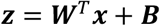

where ***x*** = *x*_1_, …, *x*_*f*_ is the usual vector of factors, ***W*** is a *f* × *n* matrix of weights, and ***B*** = *B*_1_, *B*_2_, …, *B*_*n*_ is a vector of biases, one for each possible choice. Then, one applies a smooth version of the argmax function, called *softmax*:

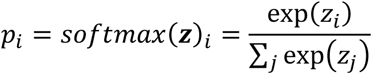

For example, if there are 3 possible choices, and we assume for simplicity that *z*_3_ = 0, then the probability *p*_1_ of the 1^st^ choice is largest when *z*_1_ > 0 and *z*_1_ > *z*_2_, dividing the space of probabilities in three regions where the three choices are more probable than the others (Figure 8**a**).

**Figure 8.**
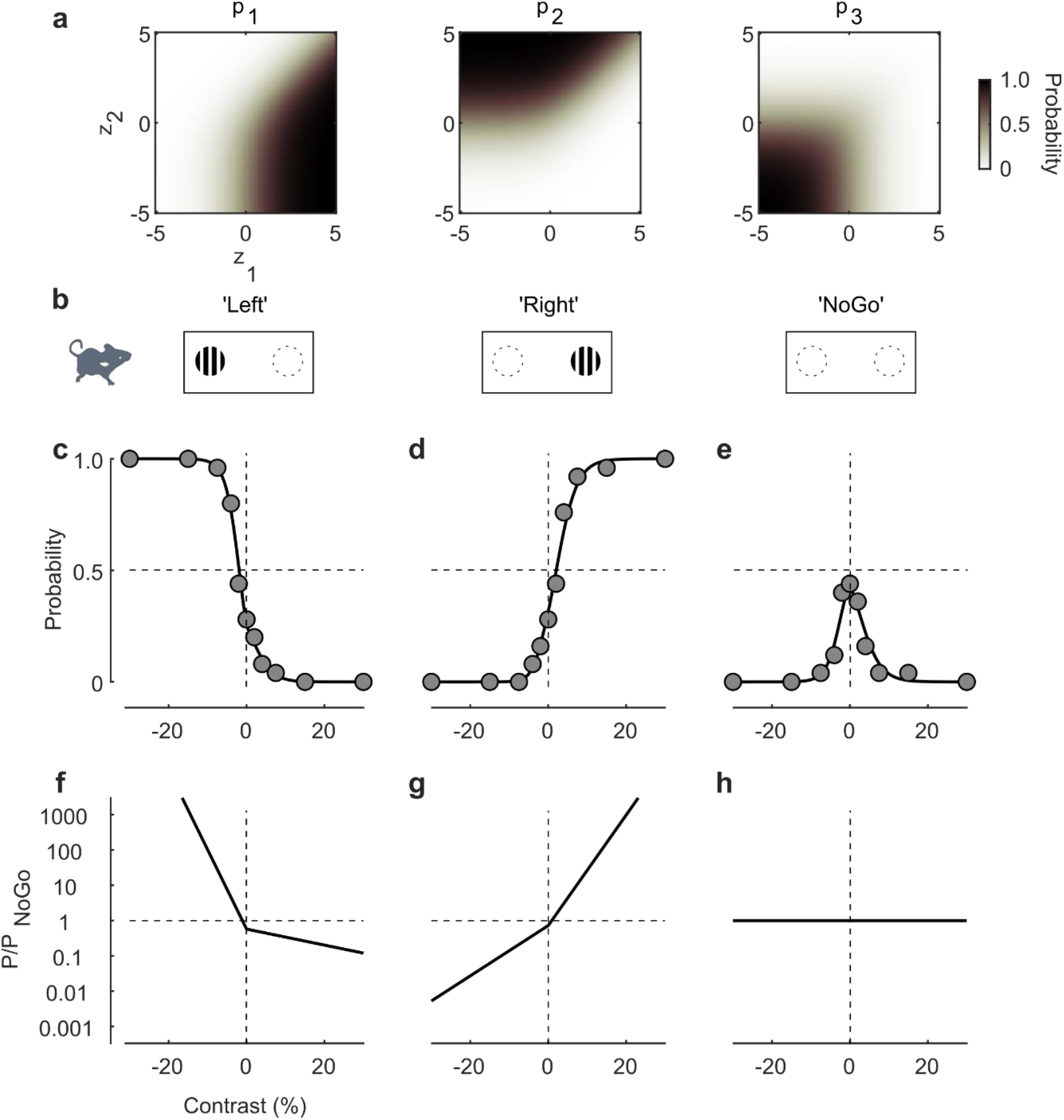
Logistic classification with more than two choices. **a**. Multinomial logistic classification among 3 choices, showing probability of choices 1,2, and 3 (left to right) as a function of z1 (abscissa) and z2 (ordinate), when z3 = 0. **b**. The 3-choice version of the position detection task, where mice choose “Left” or “Right” to indicate stimulus position, and “NoGo” if the stimulus is absent. **c-e**. Probabilities of the three choices for a mouse in this task (Mouse I from Fig. 3 of Ref.^24^), as a function of stimulus contrast. **f**-**h**. The odds of the three choices relative to the third choice.

Reassuringly, when there are only two outcomes the softmax function reduces to the familiar logistic sigmoid *p* = *σ*(*z*) with decision variable *z* = *z*_2_ − *z*_1_:

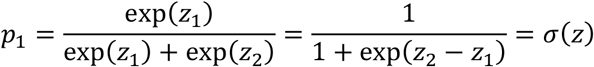

In a network, the softmax can be thought of as a competitive process among units that each receive one *z*_*i*_ as external input^5^. Because the outputs of softmax sum to 1, an increase in the input to one unit decreases the activity of others. When the difference between the largest *z*_*i*_ and the others is large, this process becomes a winner-take-all.

A logistic classifier with multiple outcomes can provide excellent fits to psychophysical data^24,25^. For example, consider a 3-choice version of the position detection task^24^, where the mouse chooses the stimulus position as “Left” or “Right” if the stimulus is present, and chooses “NoGo” if it is absent (Figure 8**b**). This three-choice design was originally devised in the framework of signal detection theory^81,82^ but it is particularly easy to analyze in terms of logistic classification, where it boils down to two equations^24^:

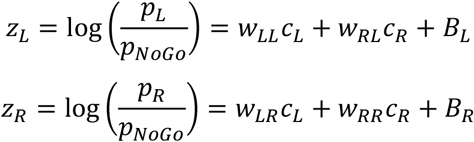

where *c*_*L*_ and *c*_*R*_ are the stimulus contrasts on the left and right. These equations provide good fits to the mouse choices (Figure 8**c**-**e**). As usual, the model is easiest to understand in terms of log odds, i.e., by plotting the values of *z*_*L*_ and *z*_*R*_ (Figure 8**f**-**h**).

## Discussion

A growing body of evidence indicates that the brain uses logistic classification to make choices based on a multitude of factors alone or in combination. These factors include relative value, relative probability, sensory inputs from one or more modalities, recent history, and brain manipulations.

To illustrate this body of evidence, I have followed a hybrid approach, in some cases reviewing existing results and in others applying the logistic classification model to data that were previously analyzed with other approaches. I acquired most of these data manually by clicking on the data points in the published plots. This introduced noise, and is particularly an issue when the probabilities are near 0 or 1, where small variations have large effects on the log odds. Given these limitations, the fits that I provided are evidence for plausibility, but not for superiority over other fits.

I have shown that an observer that performs logistic classification, and that believes that the factors are at least potentially informative, should set its bias to perform “probability matching” (or “value matching”), because this will maximize the reward. Indeed, as we have seen, observers typically adjust their bias to approximate this optimum value. If there are no informative factors, these strategies are suboptimal: it would be better to always choose the option that is more likely or has higher value (Section S3).

This suggests a reason for probability matching and value matching: the observer does not know that the evidence is irrelevant or absent (i.e. that they are gambling). Rather, they act as if they were performing a task based on evidence that is at least potentially informative. Indeed, we have seen that logistic classification is also widely used in cases where it is not optimal, for instance when stimuli are absent (and choices exhibit probability matching or value matching) or stimuli of two modalities are not independent. This suggests that logistic classification is a heuristic that the brain implements in a variety of conditions even when it is not strictly optimal^65^.

Throughout the paper, I have described logistic classification as an addition of factors that each contribute to the log odds of a choice. However, we could have just as easily talked about multiplication of factors, each of which contributes to the odds, and we could have taken the logarithm only after the multiplication. Mathematically, the two are identical but numerically it is easier to deal with log odds that vary, say, between -5 and 5 (for a natural logarithm), than with the corresponding odds, which would vary between less than one hundredth and over a hundred. Using firing rates, it is easier to encode the former than the latter. Indeed, when decoding the activity of neurons in prefrontal cortex the activity seems more commensurate to log odds^13,23^.

As to how the brain may perform logistic classification, I presented two implementations that correspond to well-established models. One echoes the framework of signal detection theory^27-29^, as it involves taking the difference between two stochastic variables and thresholding the result. The other is the well-established drift diffusion model^2,30-32^, which performs logistic classification if the starting point is midway between the bounds. Logistic classification, therefore, bridges two established models of decision making: signal detection theory and drift diffusion.

Finally, we have seen that logistic classification is a good model not only for how the brain makes decisions, but also for how these decisions are modified by manipulations such as local brain inactivations. Indeed, the effects of inactivations can often be summarized by changes in a few weights, suggesting that the inactivating regions are necessary for processing the corresponding factors.

Going forward, a promising direction is to develop the dynamic aspect of logistic classification. For instance, logistic classification can capture the effect of stimulus fluctuations, teasing apart the dynamics of cognitive computations^83,84^. At a longer time scale, it seems clear that the weights of the logistic classifier should not be considered constant, because they change with factors such as learning and brain state^51,52^.

Another important goal is to understand what limits the weights assigned to sensory stimuli, and therefore what limits performance. In some sensory tasks, the weights are limited by sensory noise (as typically captured by signal detection theory, Section S1). However, in many other tasks, the limiting factor is the appropriateness of the decoder. For instance, in an orientation discrimination task, the limiting factor is not the ability of the primary visual cortex to represent orientation, which is spectacularly high even in mice^85^, but the ability of the rest of the brain to decode that representation as required by the task. Another potential limiting factor is a general hesitance to assign large weights to sensory stimuli.

Finally, a promising direction is to use neural data as a predictor. Ultimately, a model of choice should be based on neural data, and a way to start on this path is to add neural activity to task factors^76^ such as those considered in this paper. Ultimately, the neural activity should be superior and make any other factor irrelevant^7,86^, because it is the sole determinant of the brain’s choices.

## Acknowledgments

I thank my lab colleagues Celian Bimbard, Pip Coen, Tim Sit, Valentin Schmutz, Max Shinn, and Flora Takacs for advice and help in obtaining data. I thank Kenneth D Harris for numerous insights and for suggesting the proof in Section S2, and Josh Gold for helpful suggestions. This work was funded by the Wellcome Trust (223144), by UKRI (EP/X022366/1), and by the NIH (U19NS123716). Earlier work was supported by the GlaxoSmithKline / Fight for Sight Chair in Visual Neuroscience.

## License

This preprint bears a CC-BY-NC-ND license.

## Methods

Most of the data were taken from the original papers using a manual data grabber (the MATLAB function *grabit*.*m*). The data thus obtained are somewhat noisy, and the interested reader should refer to the original papers for the definitive data. In a few cases (noted in the legends of the relevant figures), I accessed the original data.

I then fitted these data with the 2-choice logistic classification model using the MATLAB function *glmfit*. In the case of 3 choices, instead, I used the function *mnrfit*.

To describe sensory choices influenced by priors (Figure 2 **a-f**) I fit this equation:

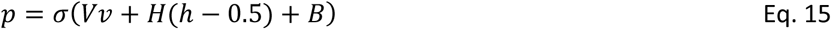

where *v* is visual strength (coherence for the motion task, contrast for the position task), *h* is the probability of a Right stimulus (determined by the block), and *V, H, B* are parameters (visual gain, prior gain, and bias).

To describe sensory choices influenced by value in humans, monkeys, and mice (Figure 2**g-o**) I fit this equation:

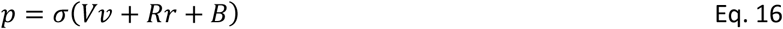

where *v* is visual strength (position, coherence, or contrast, depending on the experiment), *r* is the relative reward offered for Left vs. Right choices (determined by the block) and *V, R, B* are parameters (visual gain, reward gain, and bias).

To describe audiovisual position choices in mice (Figure 3) and visuotactile orientation choices in rats (Supp Figure 2) I fit Eq. 9. To find the best values of the exponent, I explored a range of exponents between 0.1 and 2.0, and for each value in this range I fitted a logistic classification model applied to contrast raised to that exponent. I then selected the exponent that provided the best fits.

To describe audiovisual rate choices in humans (Figure 4**a**-**h**) I fit Eq. 11.

To describe vestibulo-visual heading choices in monkeys (Figure 4**i**-**p**) I fit the same equation but with an exponent *q* applied to both *ϕ*_1_ and *ϕ*_2_.

To illustrate the effects of stimulus history (Figure 5) I simulated the model in Eq. 12 using the parameters W = 0.8, S = 2, F = -1, B = 0.

To describe the effects of brain manipulations (Figure 6) I modified Eq. 14 to allow not only for different sensitivities on L vs R but also for possible effects of manipulations:

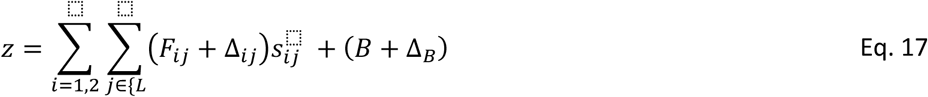

where the variables *s*_*ij*_ are the stimuli, indexed by their modality (*i* = 1 for visual contrast raised to exponent *q* and *i* = 2 for auditory position), and by their position (*j* = *L* or *R*). The parameters *F*_1*L*_, *F*_2*L*_, *F*_1*R*_, *F*_2*R*_ are the corresponding weights, and the five Δ parameters allow for the effects of the brain manipulation on the four weights *F* and on the bias *B*.

## Supplementary Figures and Mathematical Appendices

**Supp Figure 1.**
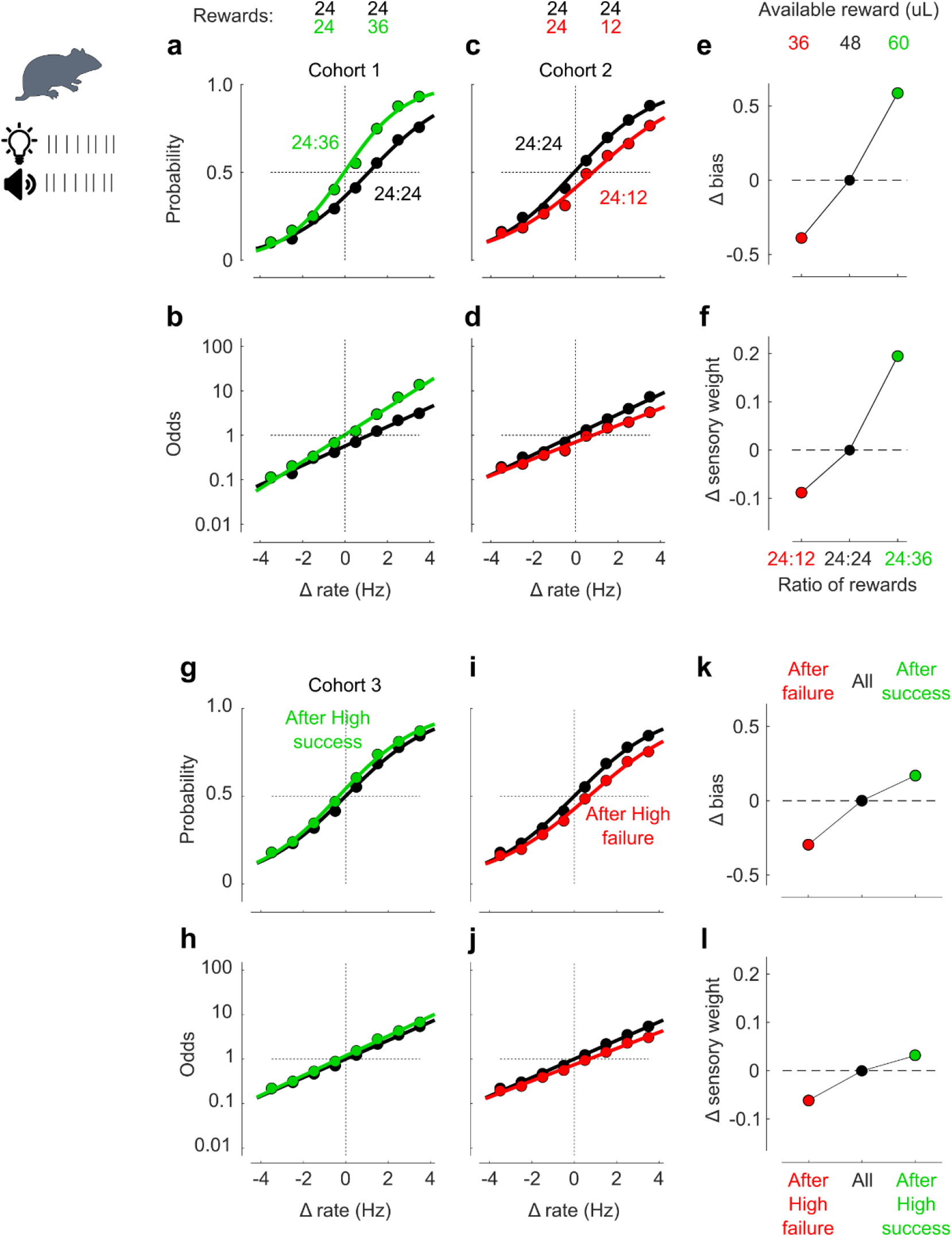
Logistic classification describes the effects of unequal rewards and of recent history. Related to Figs. 2 and 5. **a**. Probability of choosing “high” in a task where rats judge the rate of audiovisual stimuli, where the rewards for correct “high” choices are the same as correct “low” choices (24 µL, black) or higher (36 µL, green). Curves show fits of the logistic classification model where both the sensory weight W and the bias B are allowed to change when the rewards are unequal. **b**. Same data and curves, plotted in terms of the logarithmic odds of choosing “high”. **c**,**d**. Same, for a cohort of rats that were run in conditions where the rewards for correct “high” choices are the same as correct “low” choices (24 µL, black) or lower (12 µL, green). **e**. As predicted by logistic classification under the hypothesis that observers seek to maximize reward, the bias parameter was affected in opposite directions by the two manipulations. The changes in bias predicted by the model (Eq. 7) would be log(12/24)=-0.69 and log(36/24)=0.41. **f**. In addition, the sensory weight increased when the ratio of rewards increased, perhaps because the total available reward also increased (abscissa). This may have caused an increase in attention to the task variable. **g-j**. Same as a-d for a cohort of rats that were run with equal rewards for “low” and “high” correct choices, distinguishing performance following a “high” choice that was correct (green) or incorrect (red). **k**. The bias term followed the predictions of logistic classification with history terms (Eq. 12) in a win-stay lose-switch strategy. **l**. The sensory weight was slightly higher after a success than after a failure, possibly because failures tended to occur during behavioral states where the rats were paying less attention to the stimulus. Data were obtained from Fig. 4 of Ref ^26^.

**Supp Figure 2.**
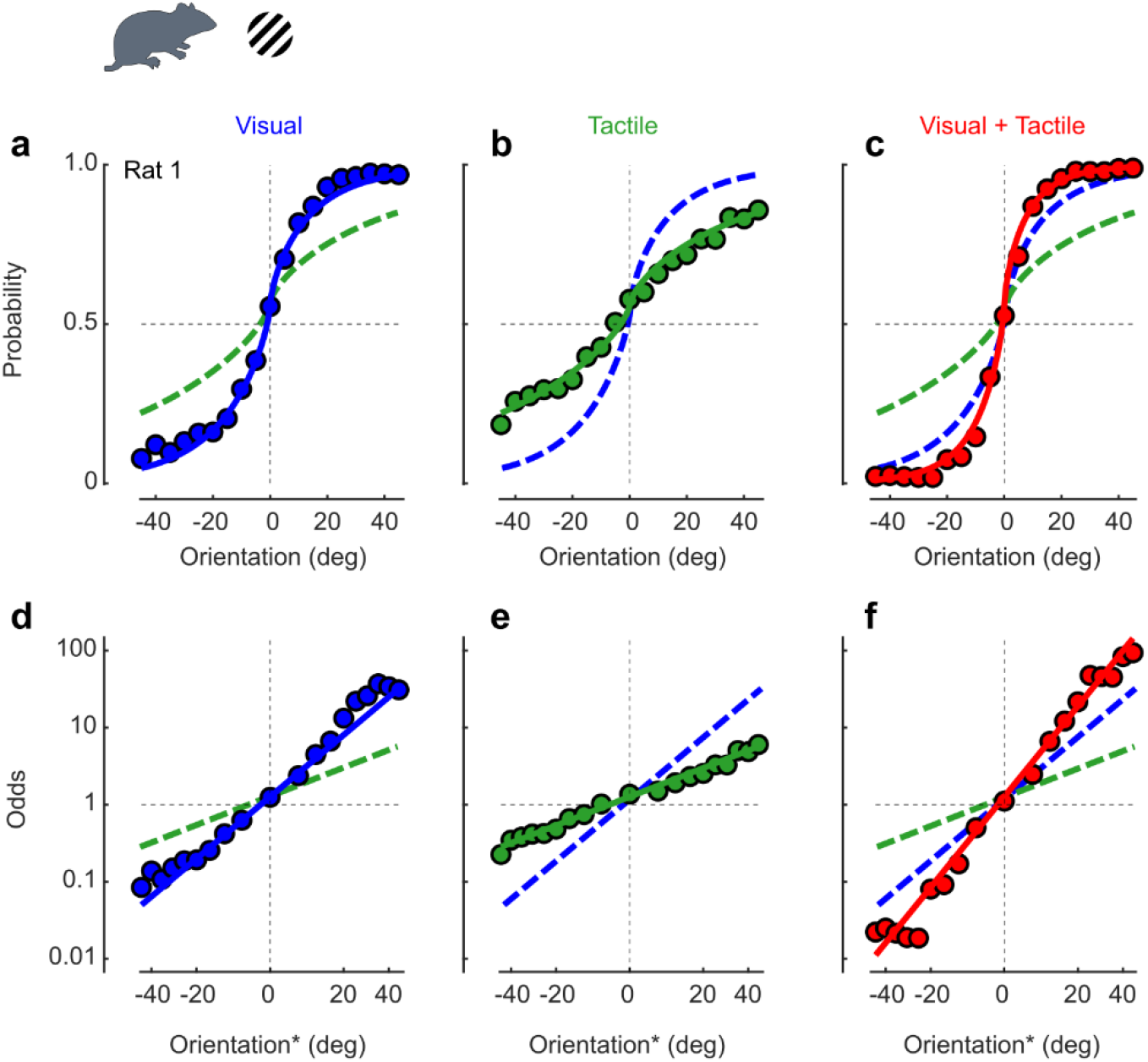
Logistic classification describes visuotactile orientation choices in rats. Related to Fig 3. **a**. Probability of Right choices for a rat performing the orientation discrimination task based on visual cues. **b**. Same, based on tactile cues. **c**. Same, based on stimuli that are both visual and tactile. **d-e**. Same data, plotted as odds of Right vs. Left in logarithmic scale. The quantity on the abscissa is compressed orientation (raised to an exponent q <1). Curves are from the logistic classification model. Data are from Rat 1 in Ref.^55^.

**Supp Figure 3.**
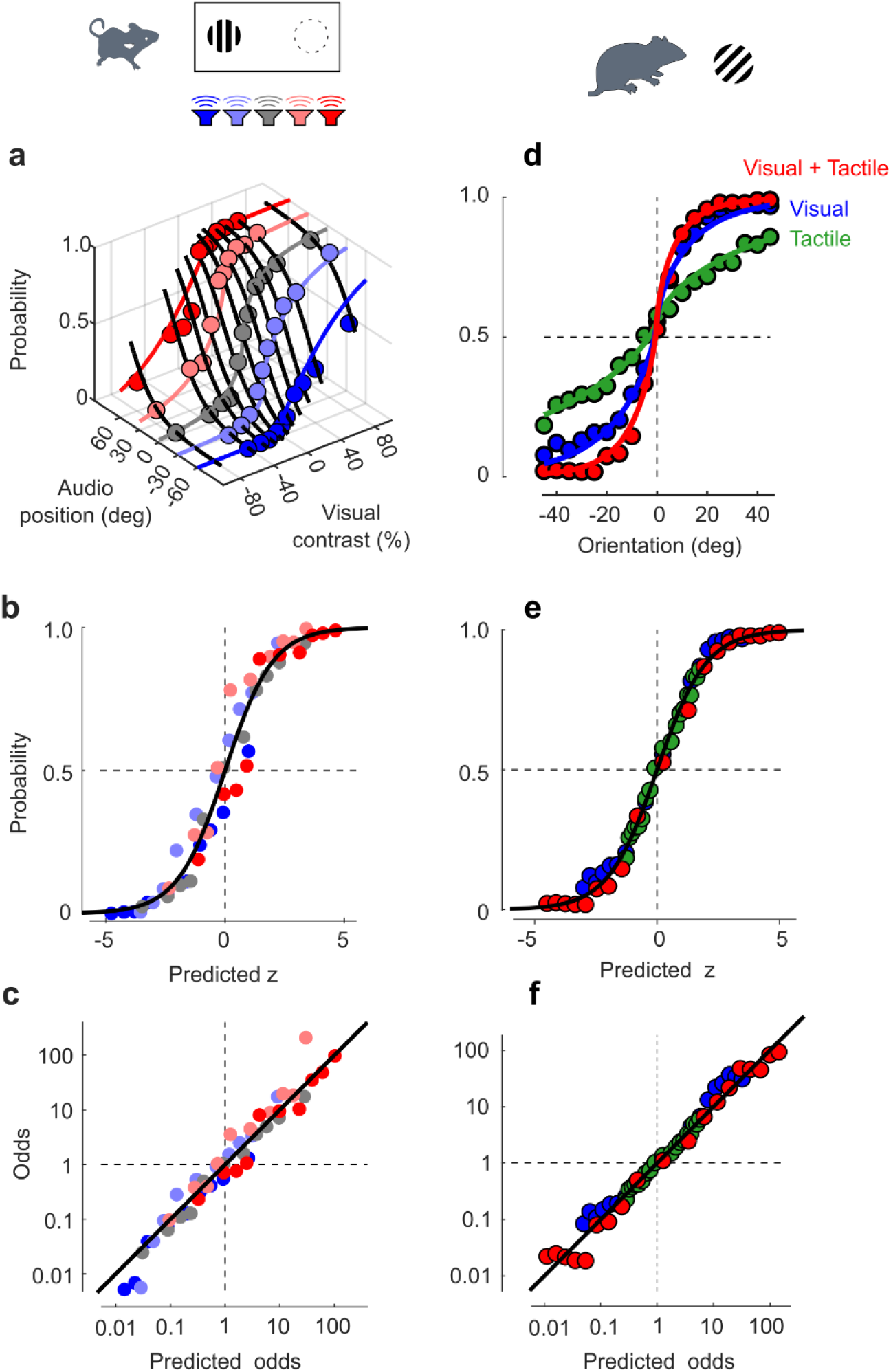
Summarizing multiple conditions on a single logistic curve. Related to Fig. 4. **a**. Probabilities of Right choices in an audiovisual task (replotted from Figure 3). **b**. The same data, plotted as a function of the decision variable z predicted by the model, plotted on top of a standard logistic curve. **c**. The same data, in log odds. **d**. Probabilities of Right choices in a visuotactile task (replotted from Supp Fig 2). Curves in a,b show fits of the logistic classification model. **e**-**f**. The same data plotted as in b,c.

### Section S1. Logistic classification and signal detection theory

We have seen that logistic classification can be obtained by taking the difference of two stochastic factors, and then making a deterministic choice by thresholding (Figure 7**b**). In this description, the utilities L, R in favor of Left and Right choices are each the sum of a deterministic value *z*_*L*_, *z*_*R*_ and a random variable *ϵ*_*L*_, *ϵ*_*R*_ (which varies across trials). The model subtracts the two and thresholds the result (Figure 9**a**). This description is mathematically equivalent to the one in the main text if the random variables are drawn from a Gumbel distribution (Figure 9**b**); in that case their difference is drawn from a logistic distribution (Figure 9**c**), and the model performs logistic classification^1,3^ (Figure 9**d**).

**Figure 9.**
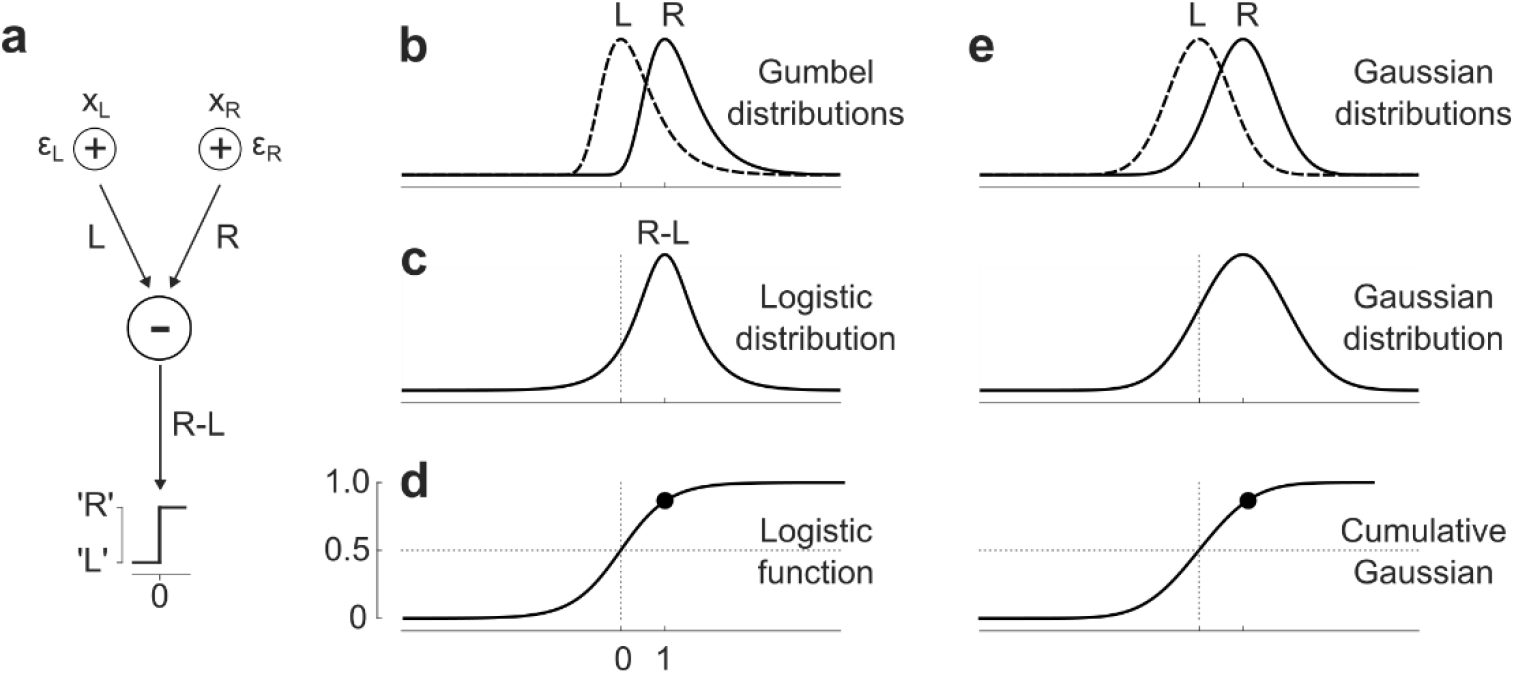
Logistic classification and signal detection theory. **a**. An alternative description of the choice model, where the utilities L, R in favor of Left and Right choices are each the sum of a deterministic value x and a random variable ε. The model subtracts the two and thresholds the result to obtain a choice. **b**. Probability distributions for the utilities L and R in the case where x_L_ = 0 and x_R_ = 1, and the random variables ε are distributed according to the Gumbel distribution. **c**. The distribution of the difference R-L is logistic. **d**. After thresholding, the probability of choosing ‘R’ as a function of x_R_ – x_L_ is a point on the logistic function. **e**. The same quantities, if the random variables ε are drawn from a Gaussian distribution. In that case the difference is Gaussian, and the probability of ‘R’ choice is a point on a cumulative Gaussian function. This is the formalism typically used in signal detection theory.

In this framework, the weight *W* that appears in the logistic classification model (Eq. 1) is the inverse of the standard deviation of *ϵ*. For an ideal observer, this standard deviation is equal to the sensory noise, but for any other observer, it is larger. The weight *W* is thus bound by the sensory reliability (the inverse of the standard deviation). It can be lower, but not higher.

This description reveals that logistic classification (for two alternatives) is similar to signal detection theory^27,28^ (SDT), which is the dominant formalism used to describe perceptual decisions. In SDT, the deterministic part of L and R is the “signal”, and the random variable is “noise”, which is thought to be sensory and related to limitations of sensory systems. In SDT, the random variable is commonly taken to be Gaussian, resulting in a psychometric curve that is a cumulative Gaussian function (Figure 9**e**).

When describing a psychometric curve determined by a single factor, the two frameworks (logistic and Gaussian) are hardly distinguishable^26,28,87-89^. For example, consider the probabilities of choosing ‘R’ in a two-alternative task (Figure 10**a**). These were generated by a logistic classification model *p* = *σ*(*Wx* + *B*), where *x* is the stimulus and *W, B* are parameters (sensory weight and bias). The same probabilities can be fitted extremely well by the cumulative Gaussian Φ((*x* − *μ*)/*σ*), where the two parameters *σ, μ* determine a “threshold” and a “bias” (Figure 10**b**).

**Figure 10.**
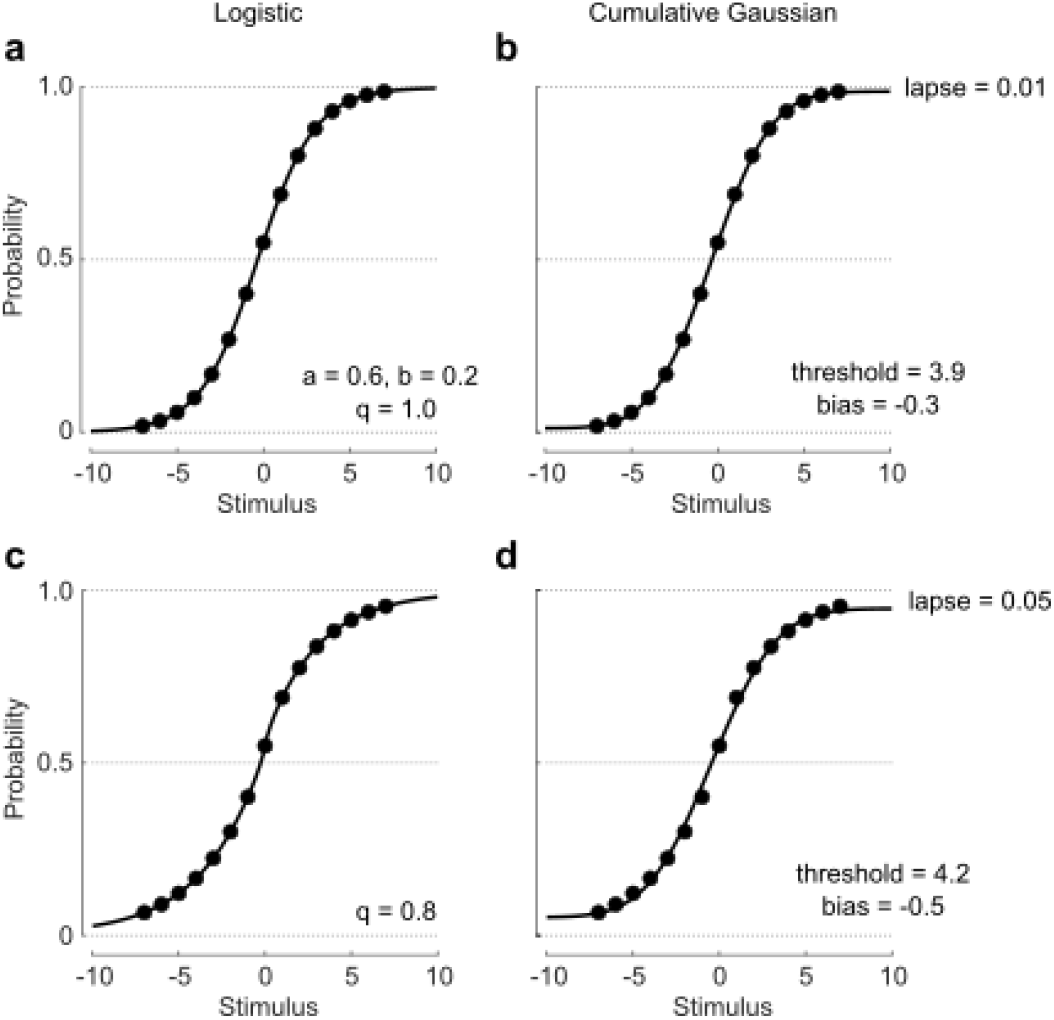
Logistic classification vs. fitting a cumulative gaussian. **a**. Probabilities simulated by a two-alternative logistic classification model (data points and curve). **b**. The same data points, fitted by a cumulative Gaussian with a lapse rate (curve). **c**. Probabilities simulated by a logistic classification model with a slight compression of the stimulus axis (exponent q=0.8). **d**. The same data points, fitted with the cumulative Gaussian with a larger lapse rate (curve).

When fitting actual data, however, the probabilities don’t always reach zero or one, even for the easiest stimuli. To deal with this eventuality, when fitting a cumulative Gaussian one typically adds an extra parameter *λ* (or two) to account for a “lapse rate” *λ* (Figure 10**d**), to obtain

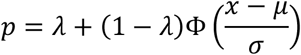

With logistic classification, it is generally not necessary to add such a lapse rate. In many cases, one can use the logistic function as is, because it tends to zero or one more gradually than the cumulative Gaussian (the logistic distribution has long tails). The probabilities are then below 1 or above 0 simply because the observer does not give enough weight *W* to the sensory input *x*. In other cases, as shown in multiple occasions in the main text, the observer may process stimulus strength in a nonlinear fashion. For instance, the sensory input *x* may be given by the measure *s* of stimulus strength used in the laboratory, raised to the power *q* < 1 to obtain a compressive nonlinearity (Figure 10**c**). In other cases, one may need a nonlinearity that is even more compressive^12,24^.

More generally, logistic classification outstrips signal detection theory (SDT) when it comes to representing more general choice environments. For instance, as I have shown in the main text, logistic classification makes it trivial to add predictors, both sensory and non-sensory, and places all these predictors on an equal footing; it has a simple linear interpretation (in terms of log odds); it makes it easy to describe the effects of brain manipulations; and it is readily extended to an arbitrary number of alternatives. Moreover, logistic classification incorporates the Bayesian assumption, without having to postulate that an observer keeps track separately of stimulus estimate and confidence (the standard deviation of the Gumbel distributions is always 1). SDT lacks some of these advantages. It can incorporate non-sensory predictors but gives them a special role^27,29^ in setting the bias *μ*. Moreover, it makes it unwieldy to have more than one or two predictors or more than two choice alternatives.

### Section S2: Logistic classification and drift diffusion

The drift diffusion model is an established model for decision making^2,30-32^ (Figure 7**c**). In the model the decision variable controls the drift rate of a random walk. When the walk hits one of the two boundaries, the corresponding decision is made.

In the general drift diffusion model, the random walk can jump at each step by an arbitrary amount, determined by a probability distribution^2,30-32,77^. In the simpler drift diffusion model described here^78^, at each moment in time the decision variable can only go up or down by 1 (Figure 7**c**).

Here I derive mathematical expressions for the probability of hitting the upper bound, and for the average time required to hit a bound. I focus on the simpler drift diffusion model (for the more general model, the derivations are more involved^31,77^).

#### Probability of hitting the upper bound

If the random walk starts from the middle between the two bounds, then the probability that it will hit one of the bounds (say, the upper one) is given by the logistic function. Specifically, here I show that given a decision variable *z* (e.g. given by Eq. 1), if the probability of going up at each step is set to *P* = *σ*(*z*/*N*), where *N* is the distance from the starting point to the bounds, the random walk reaches the upper bound with probability *p* = *σ*(*z*).

The first step to prove this is to recognize that the simplified drift diffusion model is identical to the well-known “gambler’s ruin” problem, where players 1 and 2 have *N*_1_ and *N*_2_ coins, and repeatedly bet one coin, in bets where player 1 wins with probability *P*. As shown in a textbook (Ref. ^90^, chapter XIV), the probability that player 1 will win (i.e. that player 2 will go bankrupt) is

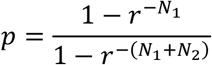

where *r* = *P*/(1 − *P*) and we have assumed that *r* ≠ 1 (the drift is preferentially in one direction).

If the two players start with equal amount of money (*N*_1_ = *N*_2_ = *N*), then player 1 will win with probability

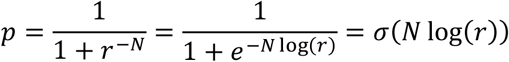

where *σ* is the logistic function. Now, if *P* = *σ*(*z*/*N*), then

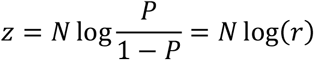

and the above expression can be rewritten as

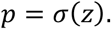

#### Expected number of steps before hitting a bound

Here I show that if the probability of going up at each step is set to *P* = *σ*(*z*/*N*), where *N* ≫ *z* is the distance from the starting point to the bounds, the drift diffusion model reaches a bound in average in *d* steps, where

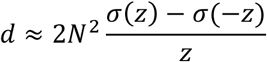

The proof starts from the same textbook (Ref. ^90^, chapter XIV), which derives the expected number of steps required for hitting a bound:

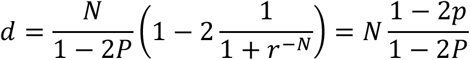

Substituting *p* = *σ*(*z*) and *P* = *σ*(*z*/*N*) we obtain:

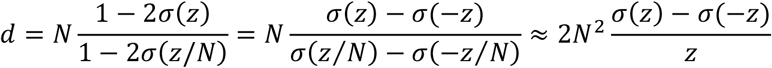

Where in the second equality I used the fact that *σ*(*x*) = 1 − *σ*(−*x*), and then I used the Taylor series approximation for *σ*(*x*) when *x* ≈ 0 (because *z* ≪ *N*) which is *σ*(*x*) ≈ 1/2 + *x*/4.

Note that the case *z* = 0 is special: the expected delay becomes *d* = *N*^2^.

#### Fitting psychometric and chronometric data

These results constitute a fundamental strength of the drift diffusion model^2,10,11,32,77^: it can account not only for how stimulus strength affects the probability of choices (the psychometric data) but also for the time of those choices (the “chronometric” data).

To see this in practice, let’s return to data measured in the random dots task^11^, where choices depend on stimulus coherence *x*, and let’s consider both the psychometric data (Figure 11**a**) and the chronometric data (Figure 11**b**). The drift diffusion model can fit the psychometric data because it provides logistic classification:

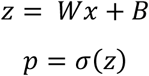

**Figure 11.**
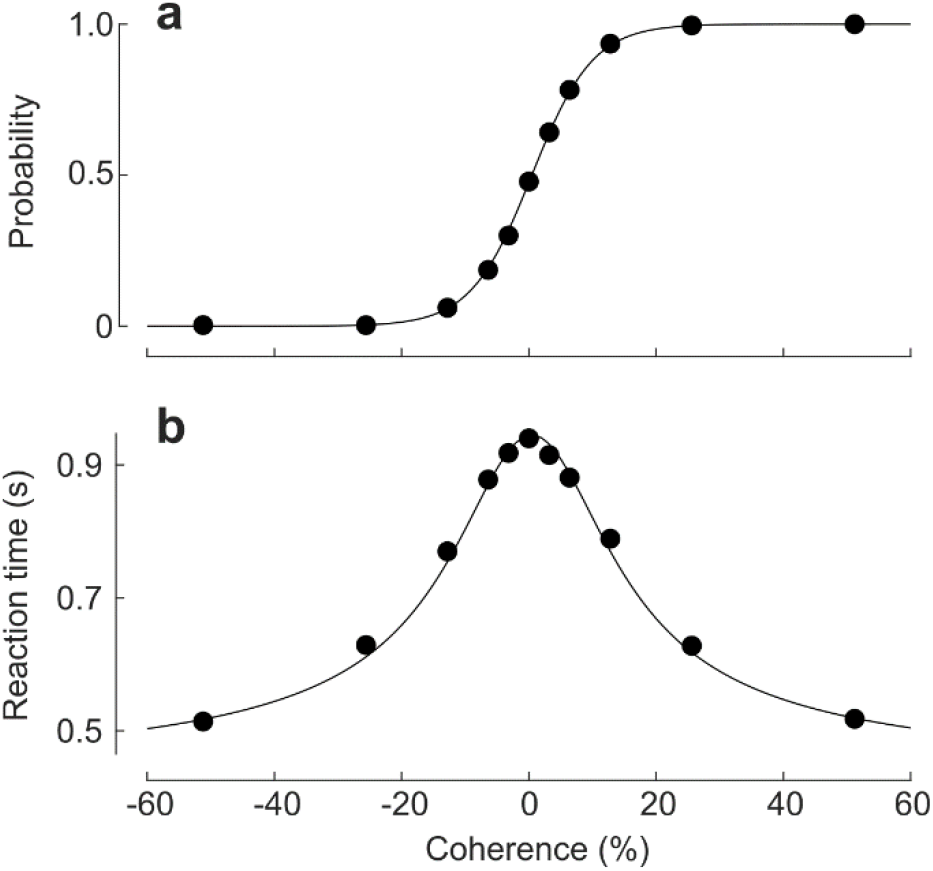
Psychometric and chronometric data fitted with the drift diffusion model. **a**. Probability of rightward choices in the random dots task, as a function of stimulus coherence (same as Figure 1d-f). I used these data to obtain the parameters W = 0.209, B = −0.097. **b**. The corresponding reaction times. I used these data to obtain the parameters R = 0.420, S = 0.525. Data were grabbed from Ref. ^11^. Curves show predictions of the drift diffusion model with the above parameters.

Moreover, it can fit chronometric data with an additional equation

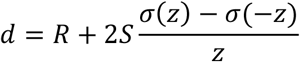

where *R, S* are parameters that describe a fixed “nondecision time” and the maximal additional time that can be added to the decision when there is no evidence. The parameters *W, B* affect both the psychometric and chronometric data. They can be fitted simultaneously to both sets of data^11^.

### Section S3: The optimal bias

Here I derive the ideal bias for an observer that performs logistic classification given two potentially different rewards for correct left vs. right choices *V*_*L*_, *V*_*R*_, and two potentially different prior probabilities of stimuli being on the left or right, *P*_*L*_, *P*_*R*_.

If an observer performs logistic classification based on evidence *x* with weight *a*, the probability that they will choose Right is *p* = *P*(′*R* ′|*x*) = *σ*(*ax* + *b*).

What is the bias *b* that maximizes expected value? The answer depends on whether the evidence *x* is informative or not.

#### Uninformative evidence

If the evidence is uninformative (*ax* = 0), the best strategy is to always choose the Right option (*b* = ∞) if *V*_*R*_ *P*_*R*_ > *V*_*L*_ *P*_*L*_ and always choose the Left option (*b* = −∞) otherwise.

This is easy to prove. The expected value of a choice is the product of the probabilities of each answer being correct (i.e. rewarded) times the corresponding reward:

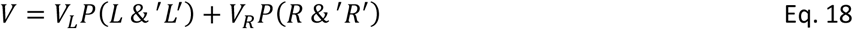

Because the choices are not guided by evidence, they are independent of the process that determines the rewarded side, the above equation can be written as

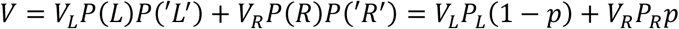

So *V* lies on a line that goes from *V*_*L*_ *P*_*L*_ to *V*_*R*_ *P*_*R*_ as *p* varies between 0 and 1. If the former value is higher than the latter, the maximum is achieved at *p* = 0, which means *b* = −∞. Otherwise, the maximum is achieved at *p* = 1, which means *b* = ∞.

#### Informative evidence

If instead the evidence is informative, the optimal bias *b* is a finite number. Specifically, if *x* is distributed uniformly (and *a* is not zero) the bias *b* that maximizes expected value is

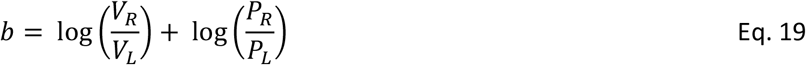

This result is proven below. My derivation is inspired by an analogous result obtained with signal detection theory using cumulative gaussians^45^. A similar expression is also obtained in the context of signal detection theory^27,29^.

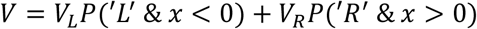

The evidence *x* is a random variable with prior probability *p*(*x*). We can thus write

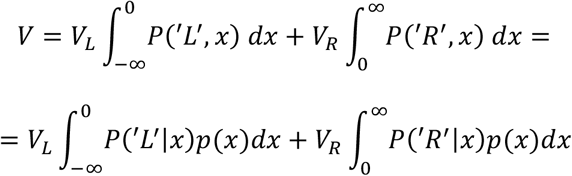

We assume that the prior probability is different between Left and Right, but otherwise flat: *p*(*x*) = *P*_*L*_ /*T, x* ∈ [−*T*, 0] and *p*(*x*) = *P*_*R*_ /*T, x* ∈ [0, *T*]. Later we will set *T* → ∞. Using the fact that *P*(′*R*^′^|*x*) = *σ*(*ax* + *b*) and *P*(′*L*^′^|*x*) = 1 − *σ*(*ax* + *b*) = *σ*(−*ax* − *b*), we obtain:

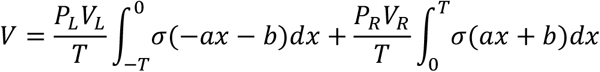

To maximize *V* we need to find *b* that such that *dV*/*db* = 0. We can bring the derivative inside the integrals:

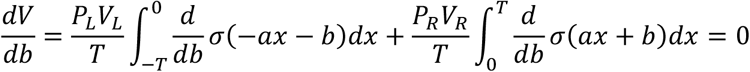

A nice property of the logistic function is that its derivative has a simple form: *dσ*(*x*)/*dx* = *σ*(*x*)*σ*(−*x*). Using this property and rearranging the terms, we obtain:

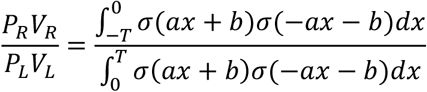

With a change of variable *z* = *ax* + *b* the Eq. becomes

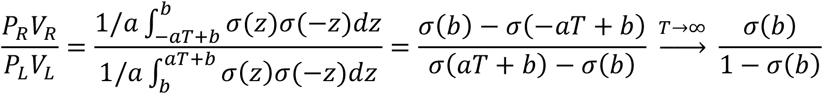

Where in the second equality we have solved the integrals by using the expression for the derivative of *σ* in reverse, and in taking the limit we have used the fact that *σ*(∞) = 1 and *σ*(−∞) = 0. If we now take the logarithm of both sides, and recognize that *σ*^−1^(*p*) = log *p*/(1 − *p*), we obtain:

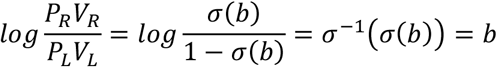

which proves the result in Eq. 19.

### Section S4: Probability matching implies logistic classification

We have seen that if an observer performs logistic classification (Eq. 5) they should set their bias to the log prior odds (Eq. 6), which leads to probability matching (when the evidence is zero).

Here I show that this argument can be inverted: the only way to achieve probability matching (for any strength of evidence) is to perform logistic classification (Eq. 5), with bias set to the log prior odds (Eq. 6).

Consider a task where a stimulus is on the Left or the Right with probabilities *P*_*L*_ and *P*_*R*_, and the observer infers the stimulus position from a sensory input *s*.

Assume that the observer aims to perform “probability matching”, i.e., to say that the stimulus is on the Right with probability *p* = *P*(*R*|*s*), the probability that the stimulus *s* is indeed on the Right.

By Bayes’ rule, the odds that the observer chooses Right are the product of two terms: one that depends on the sensory stimulus (the Bayes factor) and one that does not (the prior odds):

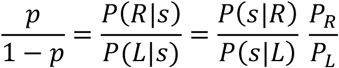

Taking the logarithm yields a sum of two terms:

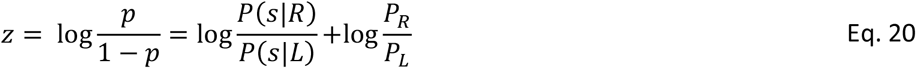

In other words, the decision variable is the sum of a function of the stimulus and a constant bias,

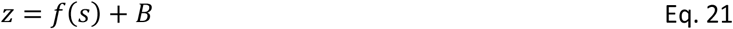

where *f*(*s*) is a (possibly nonlinear, and possibly complicated) function of the stimulus, and *B* = log(*P*_*R*_ /*P*_*L*_).

We have thus obtained the equation for logistic classification, with a suitable choice of evidence *x* = *f*(*s*) (Eq. 5), and the bias set to the log prior odds (Eq. 6).

A more general expression can be obtained if there two sensory stimuli *s*_1_, *s*_2_ of different modalities (e.g., vision and audition), and these stimuli are statistically independent^23^. In that case, using both Bayes rule and the definition of independence, *P*(*s*_1_, *s*_2_|*R*) = *P*(*s*_1_|*R*)*P*(*s*_2_|*R*), one obtains

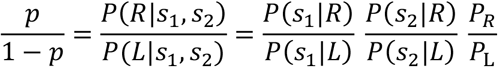

In other words,

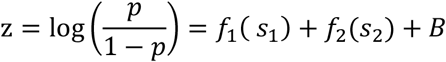

thus proving Eq. 8.

## Notes

### Competing Interest Statement

The authors have declared no competing interest.

### Summary of Updates

Made many changes. The most important one is that the paper now includes a section where logistic classification is achieved with a deterministic decision rule.

